# ABL Kinases Modulate EZH2 Phosphorylation and Signaling in Metastatic Triple Negative Breast Cancer

**DOI:** 10.1101/2025.02.18.638898

**Authors:** Ashley Colemon, Tatiana Prioleau, Clay Rouse, Ann Marie Pendergast

**Affiliations:** Department of Pharmacology and Cancer Biology, Duke University School of Medicine; Durham, NC, USA; Division of Laboratory Animal Resources, Duke University School of Medicine; Durham, NC, USA

## Abstract

Triple-negative breast cancer (TNBC) remains a leading cause of cancer associated deaths in women owing to its highly metastatic potential and limited treatment options. Recent studies have shown that expression of proteins associated with epigenetic regulation of gene expression are associated with metastatic relapse, however targeting epigenetic regulatory proteins has not resulted in effective therapies for TNBC in the clinic. The ABL tyrosine kinases promote metastasis of breast cancer cells in mouse models. However, a role of ABL kinases in the regulation of epigenetic processes in solid tumor metastasis remains unexplored. Here we show that inactivation of ABL kinases in bone metastatic TNBC cells led to a significant enrichment in gene signatures associated with the PRC2 protein complex, revealing a functional link between ABL kinases and the PRC2 complex. ABL inactivation promotes EZH2-T487 phosphorylation through the regulation of a FAK-CDK1 signaling axis. We find that phosphorylated EZH2 T487 or a phosphomimic EZH2 T487D mutant exhibit increased binding to non-canonical binding partners of EZH2 including c-MYC and ZMYND8. Notably, we identify a therapeutic vulnerability in TNBC cells whereby combination treatment with ABL allosteric inhibitors and EZH2 inhibitors elicits a synergistic decrease in TNBC cell survival in vitro, and impairs TNBC metastasis, prolonging survival of tumor-bearing mice treated with the combination therapy.

**One Sentence Summary:** ABL Kinases indirectly impact EZH2 catalytic activity by blocking a signaling cascade that leads to changes in the phosphorylation, protein interactions, and function of the PRC2 catalytic component EZH2 in TNBC.

## INTRODUCTION

Triple-negative breast cancer (TNBC) exhibits high propensity to metastasize to multiple sites including bone and brain, and metastatic TNBC is associated with high mortality rate (*1, 2*). Because TNBC tumors lack expression of targetable hormone and growth factor receptors, standard of care has been limited to chemotherapeutic options that fail to produce durable responses (*2*). Thus, patients with TNBC often experience metastatic relapse and therapy-resistance (*1, 2*). Few somatic mutations have been associated with breast cancer metastasis, but recent reports have shown that specific chromatin landscapes are linked to metastatic relapse (*3*). It was reported that therapeutic resistance of brain metastatic breast cancer cells can be induced in part by changes in the tumor chromatin landscapes mediated by interactions of breast cancer cells with stromal cells in the brain tumor microenvironment(*4*). Notably, approximately 50% of all human cancers have mutations in chromatin modifying proteins (*5*). However, single agent treatment with drugs targeting epigenetic modulators have not yet been successful, owing in part to tumor cell plasticity and therapy-resistance(*6*). Thus, new therapeutic strategies are needed to treat metastatic TNBC.

Elevated expression of ABL kinases in TNBC tumors correlates with poor survival outcomes, and promotes osteolytic metastasis in mouse models (*1, 7*). The ABL family of tyrosine kinases, ABL1 and ABL2, are activated downstream of oncogenic stimuli, as well as in response to metabolic stress, DNA damage, oxidative stress, and hypoxia among other stimuli (*8*). ABL kinases have been implicated in metastases of some solid tumors through the regulation of various transcription factor networks (*7, 9, 10*). We previously reported that ABL inactivation elicited a profound decrease in bone metastatic tumor burden and increased survival in mouse models of bone metastasis (*7*). However, the response to ABL inhibition was partial, as it did not completely eliminate metastases, suggesting that combination therapies might be exploited to treat metastatic TNBC (*7*). Interestingly, next-generation sequencing of 17,000 tumors to identify differential mutations and gene amplification changes between 3,496 primary breast tumors and 99 brain metastasis, revealed that *ABL1* mutations were more frequent in breast cancer brain metastasis than in primary breast cancer (*11*). Further, our analysis of an integrative database assembled from 22 publicly available datasets containing information on metastasis-related relapse, revealed that elevated *ABL2* expression was associated with decreased metastasis-free survival (MFS) in basal breast cancer. While global transcriptome analysis of basal breast cancer cells revealed that ABL kinase inhibition impaired the function of diverse transcriptional networks, a role for ABL kinases in the regulation of the chromatin landscapes in cancer cells through epigenetic modifications remains unexplored. Given the partial therapeutic responses of metastatic TNBC cells to an ABL-specific allosteric inhibitor, we asked whether ABL allosteric inhibitors might synergize with drugs targeting epigenetic regulators leading to improved therapies for metastatic TNBC(*7*).

Epigenetic alterations underlie the development of tumor metastasis and drug resistance (*12*). Enhanced expression and function of Enhancer of Zeste Homolog 2 (EZH2) is correlated with breast cancer metastasis (*13*). EZH2 is a histone methyl-transferase that functions as the catalytic subunit of the Polycomb Repressive Complex 2 (PRC2), which silences transcription by tri-methylation of lysine on histone H3 (H3K27me3) (*14, 15*). Interestingly, accumulating data has shown that EZH2 promotes tumorigenesis independently of the PRC2 complex through non-canonical interactions with transcription factors such as MYC and the androgen receptor (*16, 17*). Similar to ABL kinases, EZH2 promotes osteolytic metastasis of basal-like breast cancer cells, and similar to ABL inactivation, EZH2 depletion decreased the outgrowth of TNBC bone metastases in mice (*18*). Here we report that ABL kinases modulate the activity of EZH2 in metastatic TNBC cells. ABL kinases increase the canonical activity of EZH2 targeting H3K27me3, while also targeting non-canonical EZH2 signaling. Unexpectedly, we found that ABL inhibition enhances EZH2 binding to non-canonical targets linked to the accumulation of p-EZH2 T487, a post translational modification of EZH2 that promotes a shift in binding activity to non-canonical targets. Notably, we show that combination treatment with allosteric ABL inhibitors and EZH2 inhibitors elicits a synergistic decrease in TNBC cell survival in vitro, and impairs metastasis prolonging survival of tumor-bearing mice treated with the combination therapy. These data suggest that co-inhibition of ABL and EZH2 might be exploited for the treatment of metastatic TNBC patients.

## RESULTS

### ABL depletion leads to enrichment of a PRC2 gene signature and altered EZH2 activity in TNBC cells

ABL kinases are higly expressed in metastatic breast cancer and ABL-regulated gene expression profiles are implicated in driving metastatic progression (*7*). As recent reports have shown that specific chromatin landscapes are linked to metastatic relapse in TNBC (*3*), we asked whether depletion of ABL kinases might elicit changes in the expression of transcripts encoding epigenetic modifying proteins(*3*). Analysis of a previously generated RNA sequencing (RNA-seq) dataset to evaluate the transcriptomic profiles of ABL1 + ABL2 knockdown in a bone metastatic derivative of the MDA-MB-231 TNBC cell line 1833(*19*) versus control cells, revealed that one of the top modulated gene sets in the ABL double knockdown cells (AA) relative to scrambled control (SCR) corresponded to pathways involved in epigenetic regulation of gene expression (**Fig 1A**). Further interrogation of therapeutically tractable epigenetic signatures showed that target genes enriched upon the genetic ablation of the Polycome Repressive Complex 2 (PRC2) were also enriched upon ABL knockdown (**Fig 1B**). The PRC2 complex is comprised of core complex members EZH2, SUZ12, and EED and functions to repress gene expression via the trimethylation of Histone H3 at lysine 27 (H3K27me3) (*14*) (**Fig 1C**).

**Fig. 1.**
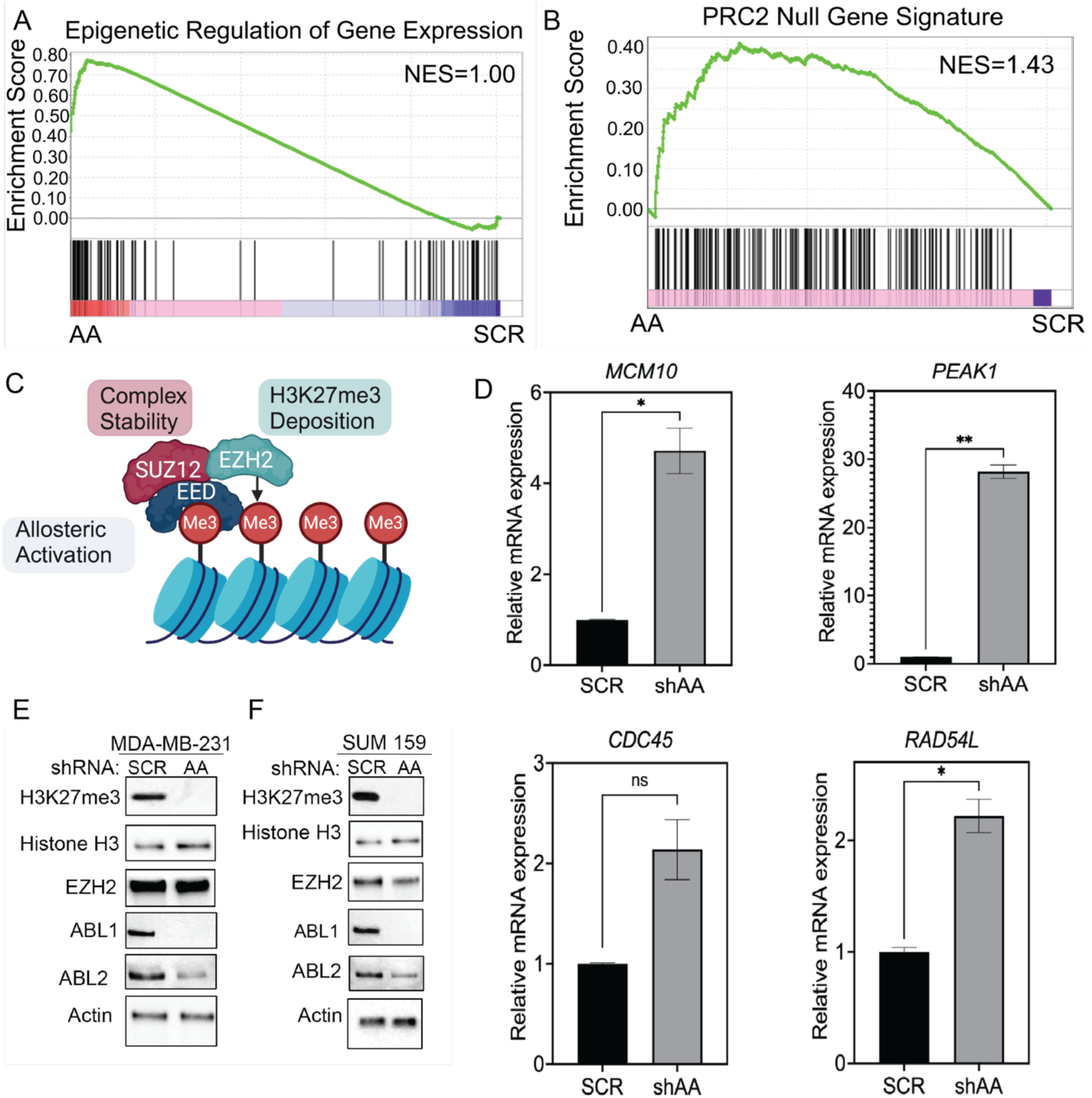
Ablation of ABL Kinases leads to an enrichment of PRC2 associated gene signatures and down regulation of PRC2 catalytic activity in TNBC. (A.) ABL kinases were knocked down using short hairpin RNAs in bone-metastatic triple-negative breast cancer cells (1833), RNA was extracted from knockdown cells followed analysis via by RNA-sequencing. Gene set enrichment analysis (GSEA) was retrospectively conducted on the generated dataset assessing changes in genes involved in epigenetic regulation of gene expression. (B.) GSEA was also performed to evaluate how genes altered by PRC2 knockdown are impacted in ABL knockdown (AA) cells. (C.)Schema of PRC2 complex. (D.) Genes significantly enriched in the retrospective RNA-seq analysis were confirmed via RT-qPCR in 1833 cells. Gene expression was normalized to a housekeeping gene (18s) before being calculated relative to SCR transduced cells. Significantly enriched genes are marked as indicated. (E.,F.) shRNAs targeting either ABL1 and ABL2 Kinase (AA) or a scramble control (SCR) were transduced into MDA-MB-231 and SUM159 TNBC cell lines to knock down both ABL Kinases. Cells lysates were analyzed by immunoblotting for changes in H3K27me3 levels. Actin was used as a loading control. All data represented in this figure represent n=3 independent experiments. Significance between SCR and shAA groups for gene expression data was determined using Welch’s T-test *,p<0.05, ns=not significant.

### ABL depletion or pharmacological inhibition leads to decreased H3K27me3 levels in TNBC cells

We found enrichment in the expression of PRC2 target genes upon ABL knockdown in the 1833 bone metastatic derivative of the MDA-MB-231 breast cancer cells. Among transcripts enriched in ABL knockdown cells were *MCM10*, *PEAK1*, *CDC45*, and *RAD54L* (**Fig. 1D**), all of which are known EZH2 target genes, as their expression was also enriched upon EZH2 knockdown in 1833 TNBC cells (**Fig S1**). Because depletion of ABL kinases elicited enrichment of EZH2 target genes in metastatic TNBC cells, we sought to assess the impact of ABL knockdown on histone trimethylation at lysine 27. We observed a profound depletion of H3K27me3 levels upon ABL knockdown without changes in the levels of histone H3 or EZH2 proteins (**Fig 1E**). This was also observed in SUM159 cells (**Fig 1F**). Additionally single knockdown of either ABL1 or ABL2 was also sufficient to deplete H3K27me3 levels (**Fig S2**). Similarly, depletion of H3K27me3 levels in MDA-MB-231 cells was observed with various ABL inhibitors including the ABL-specific allosteric inhibitors ABL001 (Asciminib) and GNF5, as well as the ABL-specific Proteolysis Targeting Chimera (PROTAC), GMB-475 (**Fig 2A**). These allosteric inhibitors target the unique myristate-binding site present in the ABL1 and ABL2 tyrosine kinases, a site which is not found in other protein kinases (*20–22*). Interestingly, treatment with GMB-475 also lead to a maginal reduction in protein levels of PRC2 core component members EZH2 and EED, suggesting that there may be intracellular proximity between ABL Kinases and these proteins.

**Fig. 2.**
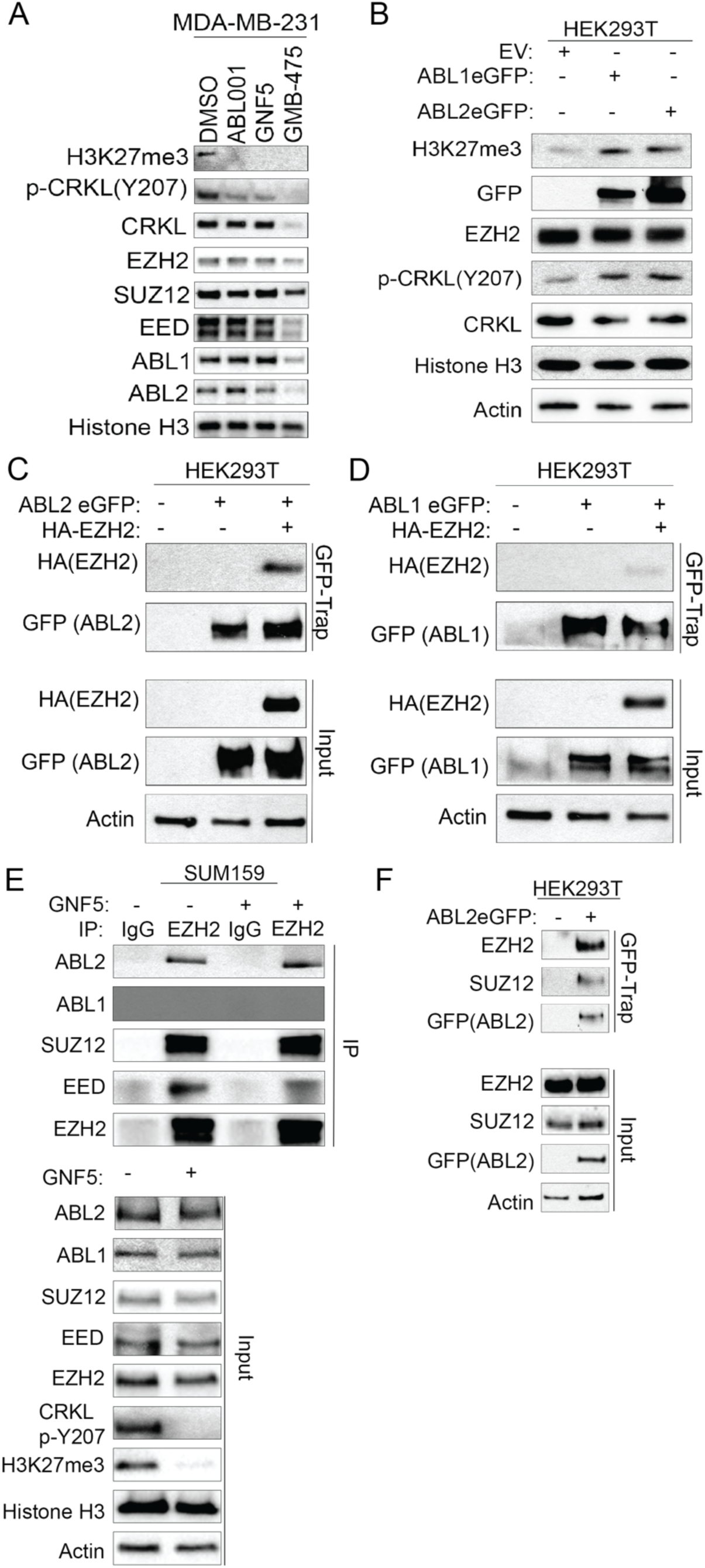
ABL Kinases activity promotes H3K27me3 accumulation and interaction with PRC2 components. (**A.**) MDA-MB-231 cells were treated with either DMSO (control), 10uM ABL001, 20uM GNF5, or 5uM GMB475 for 48 hours, cells were then assessed by immunoblotting for changes in H3K27me3 levels. Histone H3 was used as a loading control. (B.) HEK 293T cells were transiently transduced with either empty vector (EV), ABL1 eGFP, or ABL2 eGFP wildtype. Cell lysates were then assessed by immunoblotting for changes in H3K27me3 levels. Actin was used as a loading control. (C.) HEK 293T cells were transiently transfected with plasmids expressing EV, ABL2 eGFP, or ABL2eGFP and HA EZH2. ABL2 was pulled down from prepared cell lysates using the GFPtrap nanobody system. Actin was used as a loading control in the Input samples. (D.) HEK 293T cells were transiently transfected with plasmids expressing EV, ABL1 eGFP, or ABL1eGFP and HA EZH2. ABL1 was pulled down from prepared cell lysates using the GFPtrap nanobody system. Actin was used as a loading control in the Input samples. (E.) SUM159 TNBC cells were treated with 10uM GNF5 for 24 hours afterwhich endogenous EZH2 interactions were assessed via immunoprecipitation (IP) from whole cell lysate (WCL). Actin was used as a loading control in the WCL. (F.) HEK293T cells were transiently transfected with ABL2 eGFP, afterwhich ectopic ABL2 was pulled down using the GFP-Trap Nanobody system. GFP Trap pulldown lysates were probed for PRC2 complex members in tandem with WCL were analyzed by immunoblotting with indicated antibodies. Actin was used as a loading control. All data represented in this figure represent n=3 independent experiments.

### ABL kinase activity promotes H3K27me3 accumulation and interaction with EZH2

To assess the consequences of ABL kinase activation on histone K27 trimethylation, we employed a gain-of-function strategy by overexpressing wild type ABL kinases in HEK293T cells, as these cells have relatively low endogenous levels of H3K27me3 (**Fig 2B and Fig S3**). Exogenous expression of eGFP-tagged ABL1 and ABL2 wild type (WT) proteins induced a prominent enrichment of H3K27me3 levels compared to empty vector control cells (**Fig 2B**). Next, we sought to evaluate whether this H3K27me3 enrichment corresponded to association of ABL1 or ABL2 with EZH2, the primary catalytic component of the PRC2 complex. Interestingly we observed a pronounced interaction between exogenous ABL2 and EZH2, that was not observed with exogenous ABL1 and exogenous EZH2( **Fig 2C and 2D**). We then evaluated whether endogenous ABL1 and ABL2 proteins could interact with components of endogenous PRC2 complex components in the context of TNBC, and whether this interaction might be perturbed upon ABL inhibition. Pull down of endogenous EZH2 from lysates of SUM159 cells in the absence or presence of the ABL allosteric kinase inhibitor GNF5 revealed that ABL2, but not ABL1, co-immunoprecipitated with EZH2 and the EZH2-interacting SUZ12 and EED proteins (**Fig 2E**). Co-immunoprecipitation assays of eGFP-tagged ABL2 showed that not only did ABL2 interact with endogenous EZH2, but also with SUZ12, a core protein component of the PRC2 complex (**Fig 2F**). Interestingly, we found that while treatment with ABL allosteric inhibitors ABL001 and GNF5 decreased H3K27me3 levels without depleting EZH2 and associated PRC2 complex proteins in MDA-MB-231 cells, treatment with GMB475, an ABL-specific PROTAC, decreased H3K27me3 accumulation as well as the levels of endogenous EZH2, EED, and SUZ12 proteins (**Fig 2A**). This result is consistent with the recruitment of EZH2 and associated PRC2 proteins into a complex with ABL kinases, which would facilitate PROTAC-mediated degradation of closely interacting proteins within the complex upon treatment with the ABL PROTAC.

### Increased tyrosine phosphorylation signaling enhances EZH2 catalytic activity measured by H3K27me3 accumulation

To assess the role of tyrosine phosphorylation in promoting H3K27me3 levels, cells were treated with the protein tyrosine phosphatase inhibitor pervanadate to enrich for intracellular tyrosine phosphorylation (*23*). Increased tyrosine phosphorylation levels enhanced H3K27me3 accumulation in in MDA-MB-231 breast cancer cells and HEK293T cells (**Fig S3A and S3B**). To assess whether increased ABL kinase activity could enhance H3K27me3 levels we used DPH, a pharmacological activator of ABL kinases (*24*). Treatment with DPH lead to increased H3K27me3 levels in the mammary epithelial cell line MCF10A (**Fig S3C**). These data suggest that maintenance of tyrosine kinase signaling enhances EZH2 catalytic activity, consistent with reports of EZH2 phosphorylation by other tyrosine kinases (*25*).

### ABL kinase activity indirectly modulates EZH2-T487 phosphorylation and EZH2 catalytic activity in TNBC cells

Post-translational modifications of EZH2 have been implicated in changes in its activity, compartmentalization and degradation(*25–29*). To assess whether the tyrosine phosphorylation status of endogenous EZH2 was affected upon ABL kinase inhibition, breast cancer cells were treated with the ABL001 allosteric inhibitor. The tyrosine phosphorylation status of EZH2 and its associated SUZ12 protein was not significantly changed after ABL001 treatment compared to untreated controls in SUM159 breast cancer cells (**Fig 3A**). These findings are consistent with previous studies that have not detected abundant intracellular phosphorylated EZH2(*30*). These data suggest that ABL kinases may regulate EZH2 activity indirectly by modulating the activity of intermediary proteins. Analysis of publicly available clinical proteomic datasets to profile the abundance of EZH2 phosphorylation events in the context of breast cancer revealed multiple EZH2 serine and threonine phosphorylation sites (*31*). The most abundant phosphorylation site on EZH2 in-invasive-breast carcinoma clinical specimens was T487, which corresponded to approximately 20% of all detected phosphorylation events, and which was greater that phosphorylation sites attributed to AKT (S21) and p38 (T367), which only made up 4% and 2% of phosphorylation sites, respectively (**Fig 3B**). As we showed that ABL inhibition was sufficient to deplete H3K27me3 levels (**Fig 2A**), we next evaluated whether ABL inactivation could be linked to the accumulation of p-EZH2-T487 levels. Treatment with the ABL001 allosteric inhibitor resulted in marked enrichment in p-EZH2-T487 and a concomitant decrease of H3K27me3 levels indicative of decreased EZH2 catalytic activity (**Fig 3C**). A similar robust enrichment of p-EZH2-T487 in breast cancer cells was also observed in bone metastatic 1833 TNBC cells after double ABL1 + ABL2 knockdown (**Fig 3D**). Together these data support a model where inactivation of the ABL kinases indirectly promotes EZH2-T487 phosphorylation and impairs EZH2 catalytic activity as measured by decreased H3K27me3 levels.

**Fig. 3.**
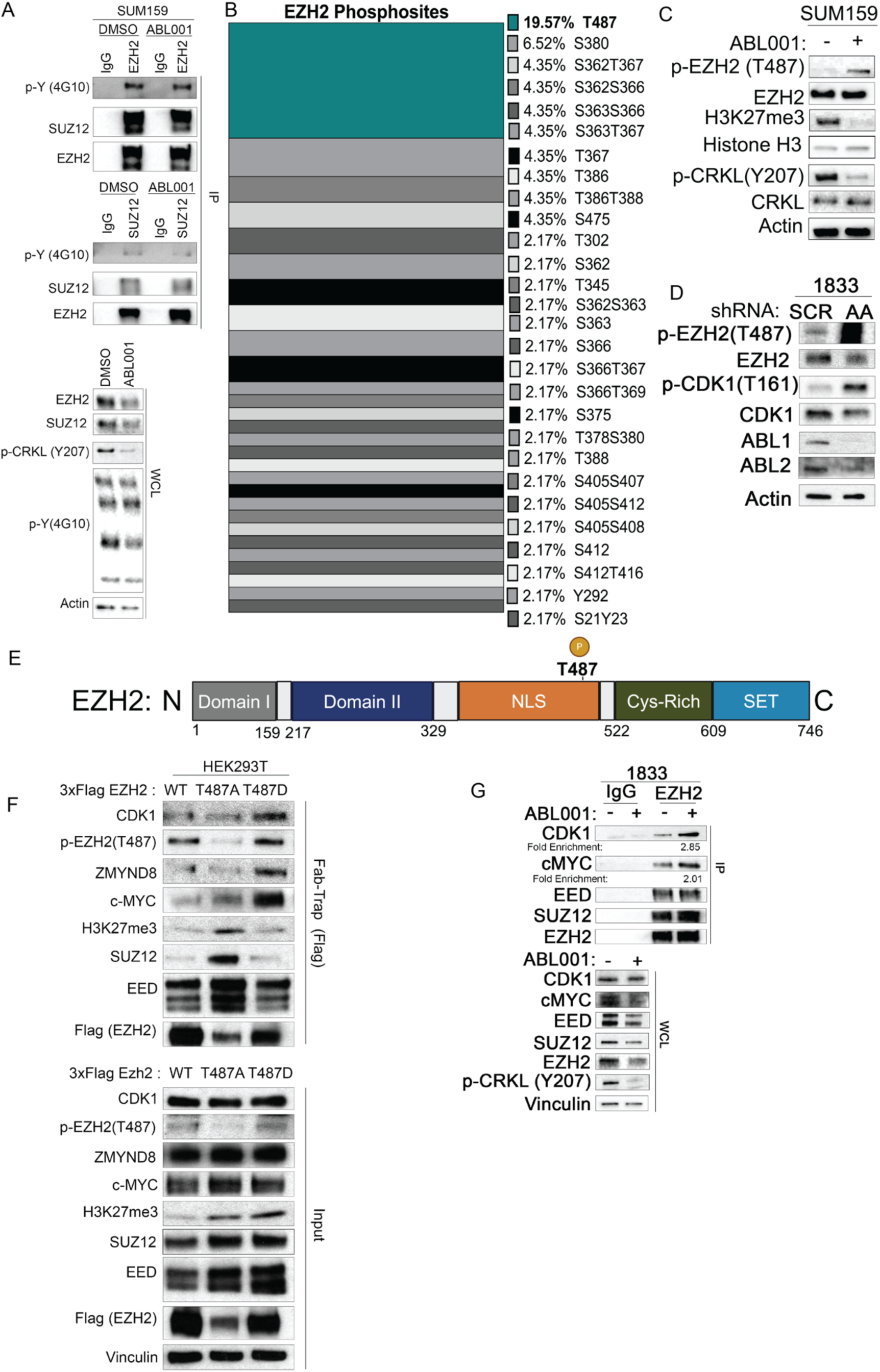
ABL Kinase activity indirectly modulates EZH2 phosphorylation in TNBC cells. (**A.**) EZH2 and SUZ12 tyrosine phosphorylation is unchanged upon ABL inhibition. SUM159 cells were treated with either DMSO (vehicle control) or ABL001 for 24 hours, after which cells were lysed and an antibody against IgG (control),EZH2,or SUZ12 was added to cell lysates. Tyrosine Phosphorylation was detected using 4G10 pan phosphotyrosine antibody. Actin was used as a loading control in the WCL. (B.) EZH2 phosphorylation mark abundance was profiled from data generated from TNBC clinical samples hosted on the Proteomic Data Commons.( C.) SUM159 cells were treated with ABL kinase inhibitor ABL001 for 24 hours with ABL001. Lysates were prepared from these cells and cells were probed for endogenous levels of phosphorylated forms of EZH2. (D.) Bone metastatic TNBC cells were lentivirally transduced with either a nontargeting shRNA (SCR) or and shRNA directed against both ABL1 and ABL2 (AA). P-EZH2 (T487) levels were assessed via immunoblot. Actin was used as a loading control. (E.) Linear protein structure of EZH2 with T487 indicated. (F.)HEK293T cells were transiently transfected with constructs expressing either pcDNA3.1_3xFlagEzh2: WT, T487A (phosphodeficient mutant), or T487D (phosphomimetic). Cell lysates were prepared and ectopically expressed EZH2 was pulled down using the fab-trap nanobody system recognizing 3xFlag. Interactions with endogenous proteins were assessed as indicated. Vinculin was used as a loading control for the input. (G.) Bone metastatic TNBC cells were treated with 10uM ABL001 for 72 hours afterwhich EZH2 was immunoprecipitated from the whole cell lysate (WCL) and probed for endogenous interactors via immunoblot. Vinculin was used as a loading control. All data represent n=3 independent experiments, with the exception of (B.).

### ABL inhibition promotes EZH2 interaction with CDK1

The phosphorylation of EZH2 at T487 is catalyzed by CDK1, and shown to decrease EZH2 methyltransferase activity via a number of proposed mechanisms including sequestering EZH2 away from core PRC2 complex members in human mesenchymal stem cells, or alternatively marking EZH2 subcellular populations for ubiquitination and degradation in HeLa cells (*30, 32*). Interestingly double ABL1 + ABL2 knockdown in 1833 cells resulted in robust induction in of active CDK1 (p-T161 CDK1) (**Fig 3D**). We next assessed whether treatment with ABL001 could enrich the interaction between CDK1 and EZH2. Indeed, pulldown of EZH2 in bone metastatic TNBC cells treated with vehicle control (DMSO) or the ABL001 allosteric inhibitor revealed that ABL inhibition enhanced EZH2 binding to CDK1(**Fig 3E**). Suggesting that ABL Kinase ablation or inhibition can stimulate phosphorylation of EZH2 on T487 in part through enriching the association between EZH2 and CDK1.

### ABL inhibition promotes altered phosphorylation of EZH2 and increases its interactions with noncanonical targets

Interactions of EZH2 with non-canonical targets have been suggested to promote tumor progression independently of its interactions with canonical targets within the PRC2 protein complex required for regulation of EZH2 histone methyltransferase activity (*17, 33*). The diverse functions of EZH2 are mediated by interactions with canonical and non-canonical protein targets, which may underlie the difficulty of targeting EZH2 activites in various tumor types (*34*). Thus, we interrogated whether ABL kinase inhibition might regulate EZH2’s interaction with its non-canonical target c-MYC (*17*). Pulldown of EZH2 in bone metastatic TNBC cells treated with vehicle control (DMSO) or the ABL001 allosteric inhibitor revealed that ABL inhibition enhanced EZH2 binding to c-MYC (**Fig 3E**). Interestingly this interaction was enriched even upon depletion of c-MYC induced by ABL kinase inhibition. In this regard, previous reports have shown that ABL inactivation decreases c-MYC expression in solid tumors including lung and meduloblastoma (*8, 35, 36*).

While EZH2 phosphorylation on T487 within the EZH2 nuclear localization sequence (NLS) domain (**Fig 3F**) has been correlated with changes in PRC2 activity, the role of this post-translational modification on EZH2 interactions with non-canonical targets is just emerging(*30, 32, 37*). In this regard, preferential association of T487 phosphorylated EZH2 with the ZMYND8 chromatin reader protein was shown in hypoxia exposed breast cancer cells and clear cell renal cell carcinoma (*38*). Using a phospho-mimic EZH2 T487D mutant, we found markedly increased association of EZH2 T487D with non-canonical targets ZMYND8 and c-MYC, while the phospho-mimic mutant protein exhibited markedly reduced association with H3K27me3, as well as reduced binding to the SUZ12 and EED components of the PRC2 complex (**Fig 3G, lane 3**). Conversely, the EZH2 phospho-deficient T487A mutant exhibited increased association with PRC2 core complex members SUZ12 and EED and also with H3K27me3. (**Fig 3G, lane 2**). The T487A mutation diminished EZH2 interaction with CDK1, while binding of the EZH2 T487D phospho-mimic mutant to CDK1 was increased compared to CDK1 binding to EZH2 WT and the T487A mutant (**Fig 3G**). These data support a model where ABL inhibition impairs EZH2 canonical function leading to decreased H3K27me3 accumulation, while promoting its interaction with non-canonical binding partners such as MYC, CDK1, and ZMYND8.

### Co-inhibition of ABL and EZH2 synergizes to selectively decrease TNBC cell viability

To assess whether co-inhibition of ABL kinase and EZH2 might be an effective therapeutic strategy in TNBC, multiple TNBC cell lines were treated with suboptimal doses of ABL kinase inhibitors, EZH2 inhibitors, or the combination of both ABL and EZH2 inhibitors, and assessed for cell survival. Combination treatment lead to a synergistic reduction in cell survival across multiple TNBC cell lines (**Fig 4A and 4B**). The synergistic decrease in cell survival is conserved across brain metastatic (MDA-BrM2a) and bone metastatic (1833) derivatives of the MDA-MB-231 cells (**Fig 4A**). Combination treatment with ABL inhibitors and EZH2 inhibitors also resulted in synergistic reduction in colony formation and decreaaed cell migration (**Fig 4C and 4D**). Combined treatment with ABL inhibitors and EZH2 inhibitors induced increased activation of Executioner Caspases 3 and 7 (**Fig S4**). Thus, combined co-inhibition of ABL and EZH2 decreases cell viability associated with increased apoptosis in TNBC cells.

**Fig. 4.**
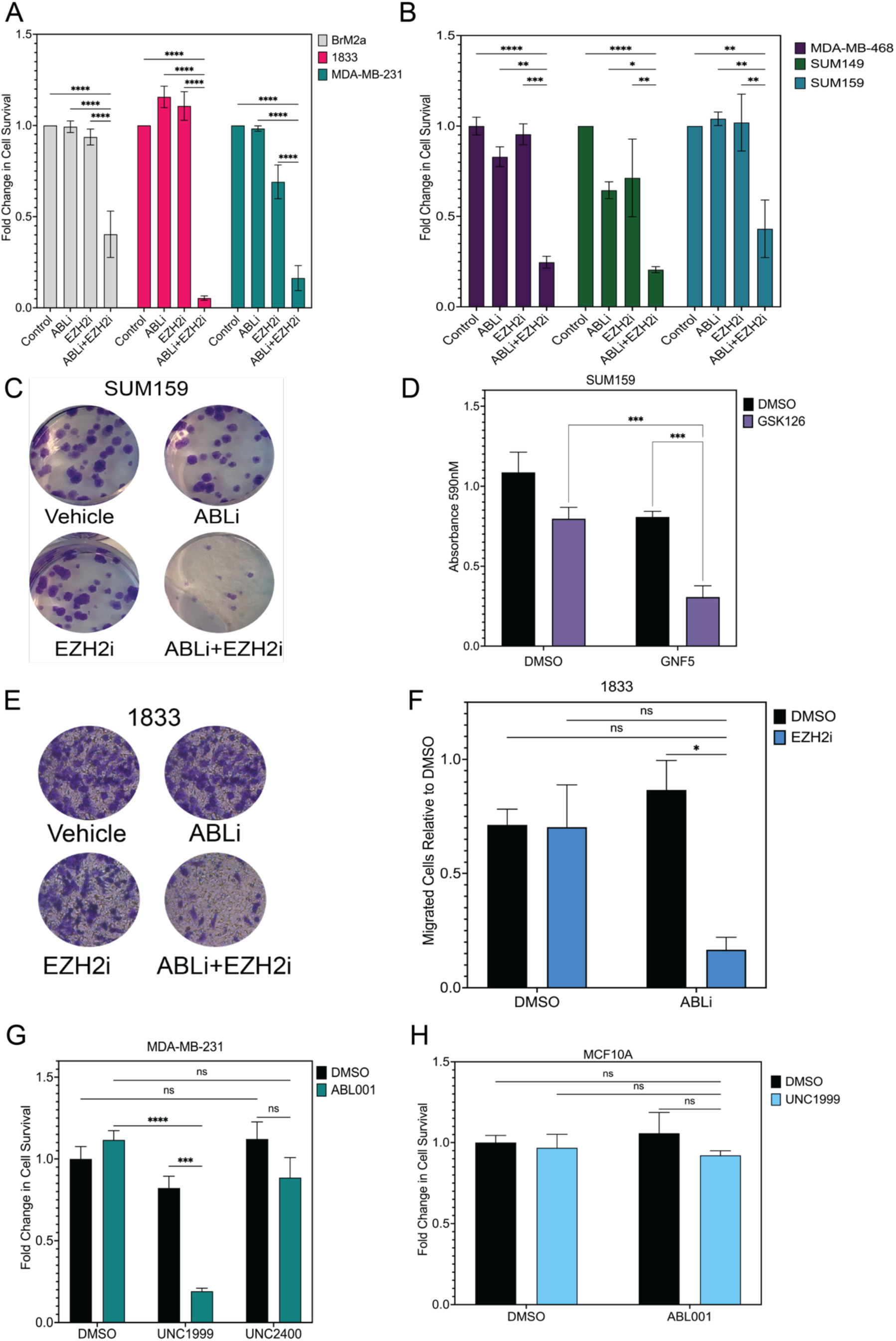
Co-inhibition of ABL and EZH2 leads to a synergistic reduction in cell viability across metastatic TNBC cell lines. (A.) TNBC cells of various origin were treated with indicated drugs at suboptimal conditions for 72 hours after which cell survival was measured using Cell Titer Glo. ABLi= 5uM ABL001 or 10uM GNF5, EZH2i= 7.5uM GSK126.(B.) TNBC cells of varying metastatic propensities were treated with indicated drugs at suboptimal conditions for 72 hours after which cell survival was measured using Cell Titer Glo. ABLi= 5uM ABL001 or 10uM GNF5, EZH2i= 7.5uM GSK126.(C,D) SUM159 cells were seeded at low density and treated with DMSO (Control), 5uM ABL001 (ABLi), 5uM GSK126 (EZH2i), or the combination of ABLi+EZH2i. Colonies were allowed to form and subsequently stained with crystal violet, and quantified as a function of staining intensity. (E.,F.) 1833 cells were seeded in Transwell Chambers and treated with DMSO, 5uM ABL001 (ABLi), 7.5uM GSK126 (EZH2i), or the combination. Cells were allowed to migrate across a serum gradient. Migrated cells were subsequently stained with crystal violet and quantified. (G.) MDA-MB-231 cells were treated with either DMSO(control), 5uM ABL001, 5uM UNC1999, 1uM UNC2400, or indicated concentrations for 48 hours afterwhich cell survival was measured using cell titer glo. (H.) MCF10A cells were treated with either DMSO, 1uM ABL001, 5uM UNC1999, or the combination of ABL001 and UNC1999 for 72 hours after which cell survival was measured using Cell Titer Glo. All data represented in this figure represent n=3 independent experiments. Significance between treatment groups was determined using a two-way analysis of variance followed by Tukey’s post hoc testing, asterisks indicate significance, *,p<0.05, ns=not significant.

EZH1 has been shown to have compensatory actions when EZH2 is inhibited, and therefore to preclude for potential compensatory effects, we employed a dual inhibitor of EZH1 and EZH2, UNC1999 (*39*). Treatment of metastatic breast cancer cells with UNC1999 lead to a reduction in cell survival as seen with GSK126 (**Fig 4E**). UNC2400 is a minimally active chemical probe that was synthesized in parallel with UNC1999 (*39*). To assess whether the synergistic decrease in cell viability was due to EZH on target action of UNC1999 we evaluated the ability of UNC2400 to synergize with ABL001. Combination treatment with ABL001 and UNC2400 did not result in a synergistic reduction in cell survival (**Fig 4E**). Furthermore, combination treatment of ABL001 and UNC1999 did not lead to a synergistic reduction in cell death of non-malignant mammary epithelial cells (**Fig 4F)**, suggesting that the combination treatment strategy may be exploited for selective synergistic killing of TNBC cancer cells without harming non-cancerous tissue.

### Co-treatment with ABL allosteric inhibitor and EZH inhibitor impairs metastatic burden and extends animal survival in mouse model of TNBC metastasis

Next, we assessed the in vivo efficacy of the combined inhibition of ABL with GNF5 and EZH1/2 with UNC1999 in a mouse model of TNBC metastasis previously employed (*40*). MDA-MB-231 1833 cells were stably transduced with a luciferase reporter before being injected intracardiac into nude athymic mice to allow for the formation of spontaneous metastasis. Seven days after implantation, mice were randomly stratified into the following treatment groups: vehicle control, 100 mg/kg GNF5, 70 mg/kg UNC1999, and combination (**Fig 5A**). GNF5 was administered once daily via oral gavage and UNC1999 was administered once daily via intraperitoneal injection. No changes in mouse body weight were observed with single and combination treatments, and tumor burden was monitored by bioluminescence imaging (BLI). Mice treated with the combination treatment exhibited the lowest tumor burden at day 27, and had the highest overall survival compared to single-treated and control mice **(Fig 5B-D)**. These data reveal that combined inhibition of ABL kinases and EZH proteins with sub-therapeutic doses of corresponding inhibitors results in enhanced overall survival in mice bearing metastatic TNBC tumor cells, revealing a potentially effective strategy to treat metastatic TNBC.

**Fig. 5.**
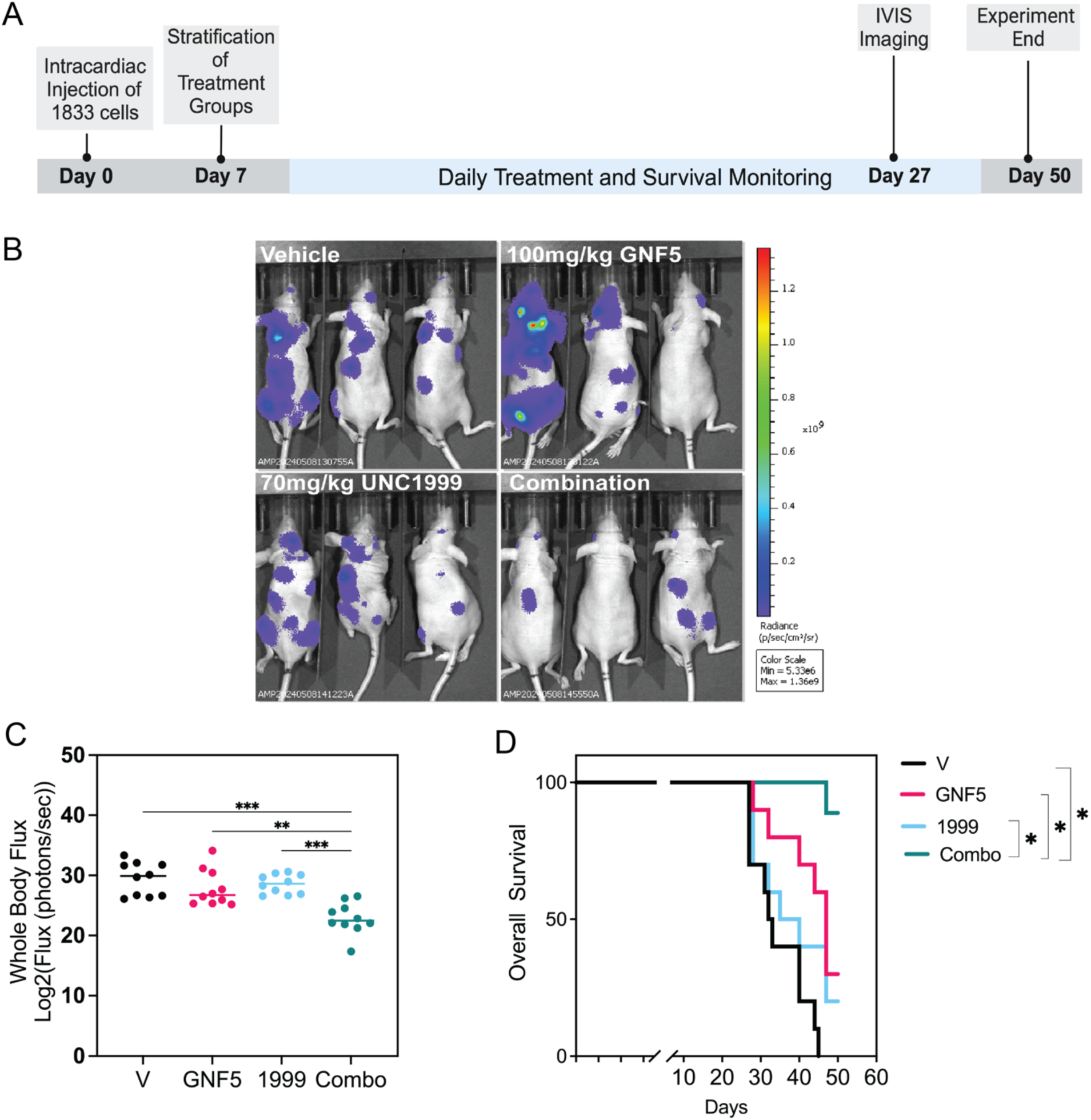
ABLi+EZH2i reduces metastatic burden and extends survival in vivo. (A) Schema of experimental design. Following 7 days post intracardiac implantation of tumor cells, mice were subjected to BLI and stratified into four treatment groups: vehicle, 100mg/kg GNF5, 70mg/kg UNC1999, or combination with 10 mice included per treatment group.(B) Representative Bioluminescent Images of mice at Day 27 post engraftment. (C) Quantification of BLI images at Day 27 post engraftment. Significance between treatment groups was determined using a two-way analysis of variance followed by Tukey’s post hoc testing, asterisks indicate significance, *,p<0.05, ns=not significant. (D) Survival analysis of treatment groups.

### ABL modulates EZH2 through a FAK-CDK1 axis

We next interrogated the mechanism by which inactivation of ABL regulates CDK1-mediated phosphorylation on T487. In silico analysis of the protein sequence of CDK1 using the Phosphosite Kinase predictor tool did not indicate any putative ABL phosphosites, but identified several Focal Adhesion Kinase (FAK) phosphorylation sites on CDK1 including Y15 and Y160 (*41*), suggesting a potential role for FAK as mediator protein linking ABL kinase activity to the regulation of CDK1 (**Table S1; Fig S5**). Functionally, Y15 on CDK1 is a key inhibitory phosphorylation site, while Y160 lies upstream of a CDK1 activation site associated with T161 phosphorylation (*42*). Similar analysis of potential sites of FAK phosphorylation by ABL kinases revealed at least 10 putative ABL phosphorylation sites, suggesting that FAK could function as an intermediary player downstream of ABL to regulate CDK1 activity in TNBC cells (**Fig S5**). The putative sites were clustered in key regulatory regions within FAK including the FERM and Kinase domain (*43*). Pulldown of ectopic FAK using the GFP-trap nanobody pulldown system in HEK293T cells, revealed that while wildtype FAK and CDK1 interact, this interaction is broken by the Y397F mutation that renders FAK kinase inactive (**Fig 6A**). We next sought to assess whether acute ABL inhibition was sufficient to induce FAK autophosphorylation, indeed we observed that treatment with allosteric ABL inhibitor ABL001 was sufficient to increase p-FAK-Y397 in SUM159 cells (**Fig 6B**).

**Fig. 6.**
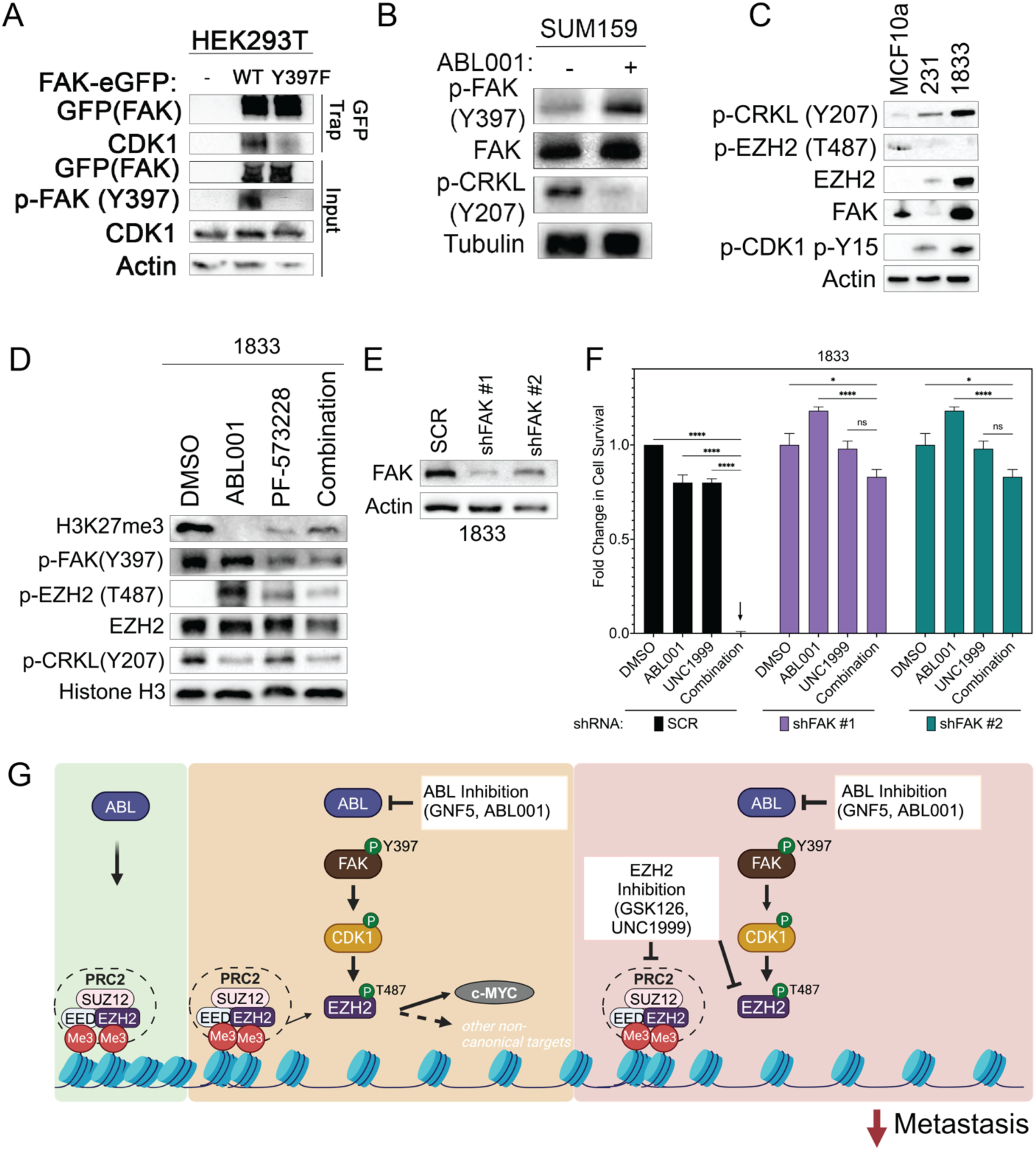
ABL Kinases regulate EZH2 through a FAK-CDK1 axis. (A.) HEK293T cells were transiently transduced with constructs expressing either EV, FAKeGFP wildtype (WT), or FAKeGFP autophosphorylation deficient mutant (Y39F). FAK was pulled down using the GFPtrap nanobody system. Actin was used as loading control in the input. (B.) SUM159 cells were treated with 10uM ABL001 for 24 hours after which cell lysates were analyzed by immunoblotting. Tubulin was used as loading control. (C.) Lysates from indicated cell lines oof increasing metastatic capacity were prepared, and levels of indicated proteins were assessed via immunoblot with Actin used as the loading control. (D.) 1833 bone metastatic TNBC cells were treated with either DMSO, 20uM ABL001, 10uM PF-573228, or the combination of both for 24 hours after which protein levels were assessed via immunoblot. (E.) 1833 bone metastatic TNBC cells were first lentivirally transduced with either nontargeting shRNA (SCR) or respective shRNAs targeting FAK (shFAK #1 or shFAK #2). Once knockdown was validated, cells were treated with either DMSO, 5uM ABL001, 7.5uM UNC1999, or the combination for 72 hours after which cell survival was measured using Cell Titer Glo. (F.) Proposed model by which ABL activity enriches EZH2 catalytic activity via inhibiting the FAK/CDK1 cascade leading to p-EZH2 T487. Conversely ABL inhibition enriches this cascade and consequently enriches p-EZH2 T487 levels. This new pool of pEZH2 rather than interacting with canonical PRC2 components is then redirected towards interactions with non canonical targets. The observed synergy observed between ABL inhibition and EZH2 manifests due to the combined effect of targeting residual PRC2 related EZH2 activity as well as interfering with the binding of EZH2 to non canonical target members. All data represented in this figure represent n=3 independent experiments. Significance between treatment groups was determined using a two-way analysis of variance followed by Tukey’s post hoc testing, asterisks indicate significance, *,p<0.05, ns=not significant.

Next, we sought to evaluate whether the FAK-CDK1 axis functions to regulate ABL-mediated phosphorylation of EZH2 p-T487 leading to phenotypic effects on the viability and metastatic phenotypes of TNBC cells. We found that increased metastatic capacity in breast cancer cells was associated with increased ABL kinase activity measured by phosphorylation of the ABL substrate p-CRKL on Y207, which was inversely correlated with p-EZH2-T487 levels and decreased CDK1 activity detected by increased phosphorylation on Y15 (**Fig 6C**). Further, the levels of FAK were highest in the bone metastatic 1833 TNBC cells (**Fig 6C**). To assess whether FAK inhibition could reverse ABL mediated regulation of EZH2 catalytic activity, 1833 TNBC cells were treated cells with either vehicle control, ABL001 allosteric inhibitor, PF-573228 FAK allosteric inhibitor, or the combination of both drugs, and evaluated EZH2 methyltransferase activity by analysis of H3K27me3 levels. Notably, co-treatment of 1833 cells with ABL001 and PF-573228 was sufficient to rescue H3K27me3 levels, and decreased p-EZH2-T487 levels back to baseline, thereby reversing the accumulation of p-EZH2-T487 induced by ABL kinase inhibition (**Fig 6D**). To ascertain that the effects of PF-573228 are on-target, we performed knockdown of FAK in bone metastatic TNBC cells with two distinct FAK shRNAs (**Fig 6E**). Depletion of FAK in metastatic 1833 breast cancer cells was sufficient to reverse the profound synergistic effects of co-inhibition of ABL (ABL001) and EZH2 (UNC1999) on cell viability (**Fig 6F**). These data are consistent with a model where ABL kinase inhibition triggers a FAK-CDK1 signaling axis leading to accumulation of p-EZH2-T487, a post-translational modification that results in a shift in binding activity to non-canonical EZH2 targets (**Fig 6G**).

## DISCUSSION

Alterations in chromatin landscapes have been linked to breast cancer metastatic relapse (*3*). Therapeutic resistance of brain metastatic breast cancer cells can be induced by changes in the tumor chromatin landscapes mediated by interactions of breast cancer cells with stromal cells in the brain tumor microenvironment (*4*). Epigenetic changes are associated with breast cancer cell plasticity and tumor heterogeneity implicated in promoting tumor progression (*44*). Further, epigenetic modulators have been implicated in driving metastatic spread through the regulation of tumor intrinsic and extrinsic signaling pathways in heterogeneous tumors such as TNBC (*4–6, 45*). Recently, it was shown that aberrant breast cancer chromatin dynamics may be regulated through contacts with distinct cell populations within the tumor microenviroment(*4*).

Canonical EZH2 activity within the PRC2 complex has been linked to the progression of several solid tumors types including TNBC (*13, 46*). EZH2 is overexpressed in 55% of invasive breast carcinomas, and has been implicated in various phenotypes of disease progression(*46*). While initial studies attributed the contribution of EZH2 to tumor progression primarily to the silencing of anti-tumor target genes as a result of H3K27me3 deposition, recent studies have also shown pleiotropic roles of EZH2’s PRC2-independent functions in tumor progression (*13, 32, 46, 47*). The PRC2-independent functions of EZH2 in TNBC have been reported to include: non-histone methylation of protein targets, transcriptional activation of pro-tumor signaling pathways, and targeting of E3 ligases to trigger protein degradation of tumor suppressor proteins(*48*). TNBC patients whose tumors express high EZH2 and low H3K27me3 have the worst prognosis upon stratification of other expression combinations, suggesting that both canonical functions and noncanonical functions of EZH2 work in tandem to promote progression of TNBC tumors (*48, 49*). Our findings reveal a role for ABL kinase activity in the regulation of canonical and non-canonical EZH2 protein complexes in TNBC cells.

The switch from canonical to non-canonical functions of EZH2 are often enabled by post translational modifications that trigger dissociation of EZH2 from other core PRC2 complex members leading to interactions with distinct targets. Phosphorylation of EZH2 at multiple sites has been implicated in switching from PRC2 dependent to PRC2 independent function (*25, 29, 32*). We found that that the tyrosine phosphorylation status of key PRC2 core complex members, including EZH2, was unchanged upon ABL inhibition. Thus, we interrogated whether ABL kinases could target signaling proteins to regulate serine/threonine phosphorylation events on EZH2. Here we identify a FAK-CDK1 signaling axis as mediator of ABL-dependent regulation of EZH2-T487 phosphorylation, complex formation, and EZH2 function in TNBC cells. Notably, FAK inhibition and knockdown rescues the inhibitory response to combined ABL and EZH2 inhibition on cell survival of metastastic TNBC cells. Further, our data show that CDK1 phosphorylation at T161 is greatly enriched upon ABL knockdown, further supporting a role for ABL-mediated regulation of a FAK-CDK1 signaling axis upstream of EZH2 T487 phosphorylation. Here we identify a FAK-CDK1 interaction that is ablated upon expression of an autophosphorylation deficient Y397F mutant of FAK.It was reported that though CDK1 is highly expressed highly in the context of breast cancer, CDK1 kinase activity is regulated by phosphorylation of key regulatory residues, some of which result in suppression of CDK1 activity (*42*).

Phosphorylation of EZH2 at T487 by CDK1 was shown to suppress PRC2 complex functions, by dissociating EZH2 from core complex members-EED and SUZ12 (*37*). Our findings reveal that treatment with ABL-specific allosteric inhibitors leads to increased EZH2 binding to non canonical targets, including MYC and concomitant increase in CDK mediated phosphorylation of EZH2 at T487. Previous studies showed that phosphorylation of EZH2 at T487 is dispensable for its intrinsic histone methyltransferase function (*30*). In contrast, we find that exogenous expression of the T487D phosphomimetic EZH2 showed decreased association of the EZH2 T487D mutant with H3K27me3 and PRC2 complex components, while this phosphomimic EZH2 mutant displayed increased association with noncanonical targets such as c-MYC and ZYMND8, the latter having recently been decribed as a preferential binding partner for p-T487 EZH2 in the context of hypoxia activated cancers (*30, 32, 38*). Our data showed that ABL inhibition or knockdown robustly induces p-T487 EZH2 leading to robust reduction of endogenous H3K27me3 levels in TNBC cells.

Some noncanonical interactions of EZH2 with various targets may contribute to cancer progression in several cancer types including TNBC (*17, 33*). The unexpected finding that ABL inhibition can enhance EZH2-T487 phosphorylation (p-EZH2-T487) promoting its binding to MYC and other targets suggest a potential vulnerability for single treatment with ABL allosteric inhibitors in TNBC cells. However, p-EZH2-T487 enrichment induced by ABL inhibition in TNBC cells doses did not significantly reduce cell survival in single treatment conditions. While, ABL kinases function to promote metastasis through engaging with transcription factor networks (*7, 9, 10, 50*), the present study reveals that ABL kinases can also support metastatic behaviors by targeting epigenetic programs.

We present data to support a model in which ABL kinases negatively engage with FAK as evidenced by enrichment of FAK activity after ABL inhibition. FAK has been implicated in regulation of EZH2 to promote TNBC bone metastasis(*18*). Here, we report that ABL kinase inhibition triggers a FAK-CDK1 signaling cascade that ultimately results in accumulation of p-EZH2-T487, a post translational modification of EZH2 that results in a shift in binding activity to non-canonical targets (*32, 38*). In this context, treatment with either an EZH2 specific inhibitor or a dual EZH1/EZH2 inhibitor blocks residual EZH2 catalytic activity leading to a reduction in metastatic burden and increased survival of tumor-bearing mice implanted with bone metastatic TNBC cells. These data suggest that co-inhibition of ABL and EZH2 might be exploited for the treatment of metastatic TNBC patients.

## MATERIALS AND METHODS

### Cell Lines and Cell Culture Methods

HEK293T, MDA-MB-231, SUM159, SUM149 cells were purchased from American Type Culture Collection (ATCC). MCF10A and MDA-MB-468 cells were purchased from the Duke Cell Culture Facility. The 1833 (bone metastatic) and BrM2a (brain metastasis) sublines were derived from the parental cell line MDA-MB-231(*51*) and were gifts from J. Massague (Memorial Sloan-Kettering Cancer Center). The HEK293T cells, MDA-MB-468, and MDA-MB-231 cells (and derived lines) were maintained in Dulbecco’s Modified Eagle’s Medium (DMEM) (Life Technologies) supplemented with 10% fetal bovine serum (FBS). The SUM149 and SUM159 cells were maintained in Ham’s F-12 Nutrient Mix (ThermoFisher) supplemented with 10% FBS, 10 mM Hepes, 5 µg/mL insulin, and 1 µg/mL hydrocortisone. MCF10A cells were maintained in MEGM™ Mammary Epithelial Cell Growth Medium (Lonza). Transient transfections were performed using Lipofectamine 2000 (Invitrogen) according to the manufacturer’s instructions. Generation of lentivirus and transductions of target cell lines were performed as described previously (*7, 52*).

For experiments assessing effects of pharmacologic agents in vitro, drugs were dissolved in dimethyl sulfoxide (DMSO) unless indicated otherwise, with the final concentration of DMSO in culture media not exceeding 0.1% vol/vol. ABL001 and GNF5 were synthesized by the Duke University Small Molecule Synthesis Facility and were validated by 1H-NMR techniques and liquid chromatography–mass spectrometry. All drugs were treated at concentrations indicated in figures and figure legends.

### Cell Survival Assay

Cell survival assays were performed as previously described with the exception of the following adjustments: Cells were seeded at a density of 1000 cells per well for all cell lines assessed(*52*). Generated results were analyzed in GraphPad.

### Cell Migration Assay

Cell migration was evaluated by plating 100,000 cells in the upper chambers of transwell chambers (8.0-μm pore size; BD Biosciences) in serum-free medium. Cells were allowed to invade for up to 48 hours in the presence of serum-containing medium in the bottom chamber. Afterward, the remaining cells and medium were removed from the upper chambers, and cells on the undersurface of the membrane were fixed, stained with crystal violet, and quantified by microscopy.

### Colony Formation Assay

Cells were seeded in 12-well plates in duplicate at 500 cells per well for SUM159s. Cells were treated with 2.5 μM ABL001, 5 μM GSK126, or combination of ABL001 and GSK126 for 1 week. Cells were then fixed with methanol and stained with crystal violet. Colonies per well were imaged, prior to destaining. To destain colonies, 1 mL of 10% acetic acid was added per well and absorbance at 590 nm was measured using a TECAN Spark microplate reader.

### Cell Lysis and Immunoblotting

Cells were lysed in RIPA buffer (50 mM Tris-HCl pH 7.4, 150 mM NaCl, 1 mM EDTA, 1% Triton X-100, 1% sodium deoxycholate and 0.1% SDS) containing protease-phosphatase inhibitor cocktail (Cell Signaling). Cell suspensions were rotated at 4°C for 10 minutes followed by microcentrifugation to remove cell debris, and protein concentration was quantified using the DC Protein Assay (BioRad). Equal amounts of protein were separated by SDS/PAGE and transferred onto nitrocellulose membranes or PVDF membranes using the Transblot Turbo Transfer system (Bio-Rad). Membranes were blocked in 5% nonfat milk in TBS-T for 1 hour prior to incubation with primary antibody overnight at 4°C. The following day primary antibodies were discarded and blots were washed 3 times with 1xTBST followed by incubation with respective HRP conjugated secondary antibody for 1 h at room temperature with gentle agitation. After incubation with secondary antibody, blots were washed an additional 3 times with TBST followed by developing using the SuperSignal™ West Pico PLUS Chemiluminescent Substrate (Invitrogen) and x-ray film or the Biorad ChemiDoc MP Imaging System.

### Real-time quantitative PCR

RNA was isolated from cells using the RNeasy RNA isolation kit (QIAGEN), and cDNA synthesis was performed using oligo(dT) primers and M-MLV reverse transcriptase (Invitrogen). RT-qPCR was performed in triplicate wells using iTaq Universal SYBR Green Supermix (Bio-Rad). Primers used in this study were purchased from Sigma Aldrich. Analysis of real-time data was collected using a Bio-Rad CFX384 machine and CFX Maestro software. The primers used in this study are listed in the supplementary materials in **Table S2**.

### Intracardiac Injections

Animal experiments were conducted in accordance with protocols approved by the Duke University Division of Laboratory Animal Resources Institutional Animal Care and Use Committee (IACUC). Intracardiac injections were performed as previously described (*7*). The ABL allosteric inhibitor GNF5 was used for *in vivo* inhibition of the ABL kinases in tumor-bearing mice and was prepared as a suspension in sterile 0.5% methylcellulose/0.5% Tween-80 as described previously (*7, 53*). The dual EZH1/EZH2 inhibitor UNC1999 was used for in vivo inhibition of PRC2 catalytic activity(*54*). GNF5 was synthesized by the Duke University Small Molecule Synthesis Facility and validated by LC-MS and 1H-NMR. UNC1999 was purchased from Selleck Chem and dissolved in sterile 0.5% methylcellulose/0.5% Tween-80 as described previously (*54*). Mice were treated with either vehicle control or 70 mg/kg/qd UNC1999 via intraperitoneal injection.

### DNA plasmids

Sequences for shRNAs used in this study are listed in the Supplementary Material in **Table S3**. EV (pN1-eGFP) and ABL2 eGFP (pN1-ABL2-eGFP WT) constructs were previously generated and described previously(*10, 50*). pcDNA3.1-3’3xFlag was a gift from Quanfu Ma (Addgene plasmid # 208616; http://n2t.net/addgene:208616; RRID:Addgene_208616); pcDNA3.1_3xFlagEzh2 WT was a gift from Thomas Cech (Addgene plasmid # 173717; http://n2t.net/addgene:173717; RRID:Addgene_173717)(*55*) pCMVHA hEZH2 was a gift from Kristian Helin (Addgene plasmid # 24230; http://n2t.net/addgene:24230; RRID:Addgene_24230)(*56*). pGFP FAK and pGFP FAK Y397F were a gift from Kenneth Yamada (Addgene plasmid # 50515; http://n2t.net/addgene:50515; RRID:Addgene_50515)(*57*).

### Site Directed Mutagenesis

The T487A and T487D EZH2 phosphomutants were cloned within the pcDNA3.1_3xFlagEzh2 WT contruct with designated primers using the Q5® Site-Directed Mutagenesis Kit (NEB) per manufacturer protocols. The primers used were as follows: T487A:F-5’-tgggaccaaagcgtgtagacagg-3’,R-5’-cctgtctacacgctttggtccca-3’;T487D:F-5’-tgggaccaaagattgtagacaggtg-3’,R-5’cacctgtctacaatctttggtccca-3’.

### Co-Immunoprecipitation

Cells were lysed in Lysis Buffer (50mM Tris pH 8.0, 150mM NaCl, 1% NP-40) with protease and phosphatase inhibitor cocktail (Cell Signaling Technology). Cell lysates were rotated at 4°C for 30 minutes, lysates were then subjected to ultracentrifugation to remove cell debris. Lysates were subsequently quantified and even protein quantities were distributed across input and IP samples. 4x SDS was added to the input samples, and they were saved for later processing. Both target directed Antibodies and Isotype IgG controls were added to respective IP conditions, and IP samples were allowed to rotate overnight at 4°C. The following day, Protein A/G beads were added to the IP samples and the samples were left to rotate at 4 °C for 3 hours. Bead immunocomplexes were pelleted with ultrafugation followed by one wash with NP-40 buffer (0.5% NP-40), and two subsequent washes with Wash Buffer (50mM Ammonium Bicarbonate). After the last wash, beads were completely dried and resuspended into 4x SDS sample buffer. Both input and IP samples were boiled and analyzed using immunoblotting.

### Nanobody Conjugated Pulldown

For GFP-Trap and Fab-Trap experiments, approaches used were similar to co immunoprecipitation approaches except in lieu of the bait antibody, nanotrap conjugated beads were added to prepared samples. Samples were allowed to incubate for 1 hour followed by washing and analysis via immunoblot.

### Gene Set Enrichment Analysis

Raw data was downloaded from the GEO via accession number GSE69125. Following analysis of RNA-seq differential expression data comparing 1833 SCR cells to 1833 shAA cells, data were sorted by statistical rank and imported into the GSEA software tool from the Broad Institute (<u>Subramanian et al., 2005</u>). Assessment of individual indicated enriched pathways were imported into the software tool and retrospectively assessed.

### In Silico Analysis of Phosphorylation Status

Clinically relevant phosphorylation statuses of proteins of interest was assessed using the National Cancer Institute Proteomic Data Commons Resource (https://pdc.cancer.gov) (*31*). Predicted target kinases for indicated phosphorylation site residues were then assessed using the Kinase Predictor Tool on PhosphositePlus ® (https://kinase-library.phosphosite.org/site) (*58, 59*).

### Antibodies

Antibodies used in this study are listed in the Supplementary Materials in **Table S4**.

### Quantification and Statistical Analysis

Statistical analyses were performed using Graphpad Prism 10 software. All data unless otherwise indicated is representative of 3 independent experiments. For qPCR data, Significance between SCR and shAA groups for gene expression data was determined using Welch’s T-test *,p<0.05, ns=not significant. Cell Survival, Cell migration, and Colony formation data represent averages ± SEM. Synergy was determined via analysis by Two-way ANOVA followed by Tukey’s post hoc testing, beinf defined as a statistically significant difference in comparison to single treatment groups. For animal experiments, Mouse numbers per group were determined through statistical power calculation based on pilot studies. Based on an expected difference of ∼30% between any two groups as indicated in pilot studies, a sample size of 10 mice per group was determined necessary to reach statistical significance based on the consideration that with a sample size of n=10, there would be a 95% likelihood that a statistically significant result would be obtained. For Kaplan-Meier survival curves, p values were calculated using log-rank (Mantel-Cox) testing. Statistical analysis of tumor flux was evaluated by ANOVA followed by Tukey’s post hoc testing to calculate p values and those less than 0.05 were quantified as statistically significant. Bar graph data represent averages ± SEM.

## Supplementary Materials

**Fig. S1.** Validation of EZH2 Target genes.

**Fig. S2.** Single ABL Knockdown ablates H3K27me3.

**Fig. S3.** Tyrosine Phosphorylation Regulates EZH2 catalytic activity.

**Fig. S4.** Combination treatment with ABL inhibitors and EZH2 inhibitors induces caspase activity.

**Fig. S5.** FAK is a putative ABL Kinase Target.

**Table S1.** CDK1 Phosphotyrosines.

**Table S2.** shRNAs

**Table S3.** RT-qPCR Primers

**Table S4.** Antibodies

## Acknowledgments

We thank the Duke University genome sequencing facility for providing assistance with deep sequencing data acquisition. We would like to thank the Duke Flow Cytometry Shared Resource for assistance with fluorescence-activated cell sorting (FACS) analysis. We would also like to acknowledge members of the Pendergast lab for their support during manuscript preparation.

## Author contributions

Conceptualization: AC, AMP

Methodology: AC,TMP, CR, AMP

Investigation: AC

Visualization: AC

Funding acquisition: AMP, AC

Supervision: AMP

Writing – original draft: AC

Writing – review & editing: AC, AMP

## Data and materials availability

All data are available in the main text or the supplementary materials.

## References and Notes

1. X. Li et al., Triple-negative breast cancer has worse overall survival and cause-specific survival than non-triple-negative breast cancer. Breast Cancer Research and Treatment 161, 279–287 (2017).

2. D. O’Reilly, M. A. Sendi, C. M. Kelly, Overview of recent advances in metastatic triple negative breast cancer. World J Clin Oncol 12, 164–182 (2021).

3. W. L. Cai et al., Specific chromatin landscapes and transcription factors couple breast cancer subtype with metastatic relapse to lung or brain. BMC Medical Genomics 13, 33 (2020).

4. M.-A. Goyette et al., Cancer–stromal cell interactions in breast cancer brain metastases induce glycocalyx-mediated resistance to HER2-targeting therapies. Proceedings of the National Academy of Sciences 121, e2322688121 (2024).

5. Jueng S. You, Peter A. Jones, Cancer Genetics and Epigenetics: Two Sides of the Same Coin? Cancer Cell 22, 9–20 (2012).

6. S. Purja, D. T. Nguyen, E. Kim, Breast cancer epigenetics: current and evolving treatment. Breast Cancer 31, 869–885 (2024).

7. J. Wang, C. Rouse, J. S. Jasper, A. M. Pendergast, ABL kinases promote breast cancer osteolytic metastasis by modulating tumor-bone interactions through TAZ and STAT5 signaling. Science Signaling 9, ra12-ra12 (2016).

8. B. Mayro et al., ABL kinases regulate the stabilization of HIF-1α and MYC through CPSF1. Proceedings of the National Academy of Sciences 120, e2210418120 (2023).

9. J. J. Gu, J. Hoj, C. Rouse, A. M. Pendergast, Mesenchymal stem cells promote metastasis through activation of an ABL-MMP9 signaling axis in lung cancer cells. PLOS ONE 15, e0241423 (2020).

10. J. P. Hoj, B. Mayro, A. M. Pendergast, A TAZ-AXL-ABL2 Feed-Forward Signaling Axis Promotes Lung Adenocarcinoma Brain Metastasis. Cell Reports 29, 3421–3434.e3428 (2019).

11. S. D. Ferguson et al., Profiles of brain metastases: Prioritization of therapeutic targets. Int J Cancer 143, 3019–3026 (2018).

12. W. A. Flavahan, E. Gaskell, B. E. Bernstein, Epigenetic plasticity and the hallmarks of cancer. Science 357, eaal2380 (2017).

13. S. H. Alford, K. Toy, S. D. Merajver, C. G. Kleer, Increased risk for distant metastasis in patients with familial early-stage breast cancer and high EZH2 expression. Breast Cancer Research and Treatment 132, 429–437 (2012).

14. Y. Shi et al., Structure of the PRC2 complex and application to drug discovery. Acta Pharmacologica Sinica 38, 963–976 (2017).

15. K. H. Kim, C. W. M. Roberts, Targeting EZH2 in cancer. Nature Medicine 22, 128–134 (2016).

16. K. Xu et al., EZH2 Oncogenic Activity in Castration-Resistant Prostate Cancer Cells Is Polycomb-Independent. Science 338, 1465–1469 (2012).

17. J. Wang et al., EZH2 noncanonically binds cMyc and p300 through a cryptic transactivation domain to mediate gene activation and promote oncogenesis. Nature Cell Biology 24, 384–399 (2022).

18. L. Zhang et al., EZH2 engages TGFβ signaling to promote breast cancer bone metastasis via integrin β1-FAK activation. Nature Communications 13, 2543 (2022).

19. J. Wang, C. Rouse, J. S. Jasper, A. M. Pendergast, ABL kinases promote breast cancer osteolytic metastasis by modulating tumor-bone interactions through TAZ and STAT5 signaling. Science Signaling 9, ra12 (2016).

20. F. J. Adrián et al., Allosteric inhibitors of Bcr-abl–dependent cell proliferation. Nature Chemical Biology 2, 95–102 (2006).

21. M. Breccia, G. Colafigli, E. Scalzulli, M. Martelli, Asciminib: an investigational agent for the treatment of chronic myeloid leukemia. Expert Opinion on Investigational Drugs 30, 803–811 (2021).

22. G. M. Burslem et al., Targeting BCR-ABL1 in Chronic Myeloid Leukemia by PROTAC-Mediated Targeted Protein Degradation. Cancer Research 79, 4744–4753 (2019).

23. J. Reiland et al., Pervanadate activation of intracellular kinases leads to tyrosine phosphorylation and shedding of syndecan-1. Biochemical Journal 319, 39–47 (1996).

24. J. Yang et al., Discovery and Characterization of a Cell-Permeable, Small-Molecule c-Abl Kinase Activator that Binds to the Myristoyl Binding Site. Chemistry & Biology 18, 177–186 (2011).

25. J. Yan et al., EZH2 phosphorylation by JAK3 mediates a switch to noncanonical function in natural killer/T-cell lymphoma. Blood 128, 948–958 (2016).

26. Z. Li et al., Post-translational modifications of EZH2 in cancer. Cell & Bioscience 10, 143 (2020).

27. T. Anwar et al., p38-mediated phosphorylation at T367 induces EZH2 cytoplasmic localization to promote breast cancer metastasis. Nature communications 9, 2801 (2018).

28. S. Chen et al., Cyclin-dependent kinases regulate epigenetic gene silencing through phosphorylation of EZH2. Nature Cell Biology 12, 1108–1114 (2010).

29. T.-L. Cha et al., Akt-Mediated Phosphorylation of EZH2 Suppresses Methylation of Lysine 27 in Histone H3. Science 310, 306–310 (2005).

30. S. C. Wu, Y. Zhang, Cyclin-dependent Kinase 1 (CDK1)-mediated Phosphorylation of Enhancer of Zeste 2 (Ezh2) Regulates Its Stability*. Journal of Biological Chemistry 286, 28511–28519 (2011).

31. Z. Wang et al., NCI Cancer Research Data Commons: Resources to Share Key Cancer Data. Cancer Research 84, 1388–1395 (2024).

32. Y. Wei et al., CDK1-dependent phosphorylation of EZH2 suppresses methylation of H3K27 and promotes osteogenic differentiation of human mesenchymal stem cells. Nature Cell Biology 13, 87–94 (2011).

33. G. J. Dardis, J. Wang, J. M. Simon, G. G. Wang, A. S. Baldwin, An EZH2-NF-κB regulatory axis drives expression of pro-oncogenic gene signatures in triple negative breast cancer. iScience 26, 107115 (2023).

34. M.-L. Eich, M. Athar, J. E. Ferguson, III, S. Varambally, EZH2-Targeted Therapies in Cancer: Hype or a Reality. Cancer Research 80, 5449–5458 (2020).

35. J. K. Jones et al., ABL1 and ABL2 promote medulloblastoma leptomeningeal dissemination. Neuro-Oncology Advances 5, vdad095 (2023).

36. J. J. Gu, et al., Inactivation of ABL kinases suppresses non–small cell lung cancer metastasis. JCI Insight 1, (2016).

37. S. Kaneko et al., Phosphorylation of the PRC2 component Ezh2 is cell cycle-regulated and up-regulates its binding to ncRNA. Genes & Development 24, 2615–2620 (2010).

38. B. Tang et al., ZMYND8 preferentially binds phosphorylated EZH2 to promote a PRC2-dependent to-independent function switch in hypoxia-inducible factor–activated cancer. Proceedings of the National Academy of Sciences 118, e2019052118 (2021).

39. K. D. Konze et al., An Orally Bioavailable Chemical Probe of the Lysine Methyltransferases EZH2 and EZH1. ACS Chemical Biology 8, 1324–1334 (2013).

40. J. Wang, C. Rouse, J. S. Jasper, A. M. Pendergast, ABL kinases promote breast cancer osteolytic metastasis by modulating tumor-bone interactions through TAZ and STAT5 signaling. Sci Signal 9, ra12 (2016).

41. P. V. Hornbeck et al., PhosphoSitePlus, 2014: mutations, PTMs and recalibrations. Nucleic Acids Research 43, D512–D520 (2014).

42. G. Massacci, L. Perfetto, F. Sacco, The Cyclin-dependent kinase 1: more than a cell cycle regulator. British Journal of Cancer 129, 1707–1716 (2023).

43. M. S. Bauer et al., Structural and mechanistic insights into mechanoactivation of focal adhesion kinase. Proceedings of the National Academy of Sciences 116, 6766–6774 (2019).

44. D. Hanahan, Hallmarks of Cancer: New Dimensions. Cancer Discovery 12, 31–46 (2022).

45. S. M. O. Phipps, W. K. Woodfin, T. O. Tollefsbol, The epigenetics of breast carcinogenesis and metastasis. Current Genomics 6, 129–135 (2005).

46. C. G. Kleer et al., EZH2 is a marker of aggressive breast cancer and promotes neoplastic transformation of breast epithelial cells. Proceedings of the National Academy of Sciences 100, 11606–11611 (2003).

47. S. Mahara et al., HIFI-α activation underlies a functional switch in the paradoxical role of Ezh2/PRC2 in breast cancer. Proceedings of the National Academy of Sciences 113, E3735–E3744 (2016).

48. T. Anwar, M. E. Gonzalez, C. G. Kleer, Noncanonical Functions of the Polycomb Group Protein EZH2 in Breast Cancer. The American Journal of Pathology 191, 774–783 (2021).

49. W. K. Bae et al., The methyltransferase EZH2 is not required for mammary cancer development, although high EZH2 and low H3K27me3 correlate with poor prognosis of ER-positive breast cancers. Molecular Carcinogenesis 54, 1172–1180 (2015).

50. J. P. Hoj, B. Mayro, A. M. Pendergast, The ABL2 kinase regulates an HSF1-dependent transcriptional program required for lung adenocarcinoma brain metastasis. Proceedings of the National Academy of Sciences 117, 33486–33495 (2020).

51. Y. Kang et al., A multigenic program mediating breast cancer metastasis to bone. Cancer Cell 3, 537–549 (2003).

52. J. H. Luttman et al., ABL allosteric inhibitors synergize with statins to enhance apoptosis of metastatic lung cancer cells. Cell Reports 37, (2021).

53. A. A. Wylie et al., The allosteric inhibitor ABL001 enables dual targeting of BCR– ABL1. Nature 543, 733–737 (2017).

54. B. Xu et al., Selective inhibition of EZH2 and EZH1 enzymatic activity by a small molecule suppresses MLL-rearranged leukemia. Blood 125, 346–357 (2015).

55. Y. Long et al., RNA is essential for PRC2 chromatin occupancy and function in human pluripotent stem cells. Nature Genetics 52, 931–938 (2020).

56. A. P. Bracken et al., EZH2 is downstream of the pRB-E2F pathway, essential for proliferation and amplified in cancer. The EMBO Journal 22, 5323–5335-5335 (2003).

57. J. Gu et al., Shc and Fak Differentially Regulate Cell Motility and Directionality Modulated by Pten. Journal of Cell Biology 146, 389–404 (1999).

58. J. L. Johnson et al., An atlas of substrate specificities for the human serine/threonine kinome. Nature 613, 759–766 (2023).

59. T. M. Yaron-Barir et al., The intrinsic substrate specificity of the human tyrosine kinome. Nature 629, 1174–1181 (2024).

