## Supplemental Materials for "ABL Kinases Modulate EZH2 Phosphorylation and Signaling in Metastatic Triple Negative Breast Cancer"

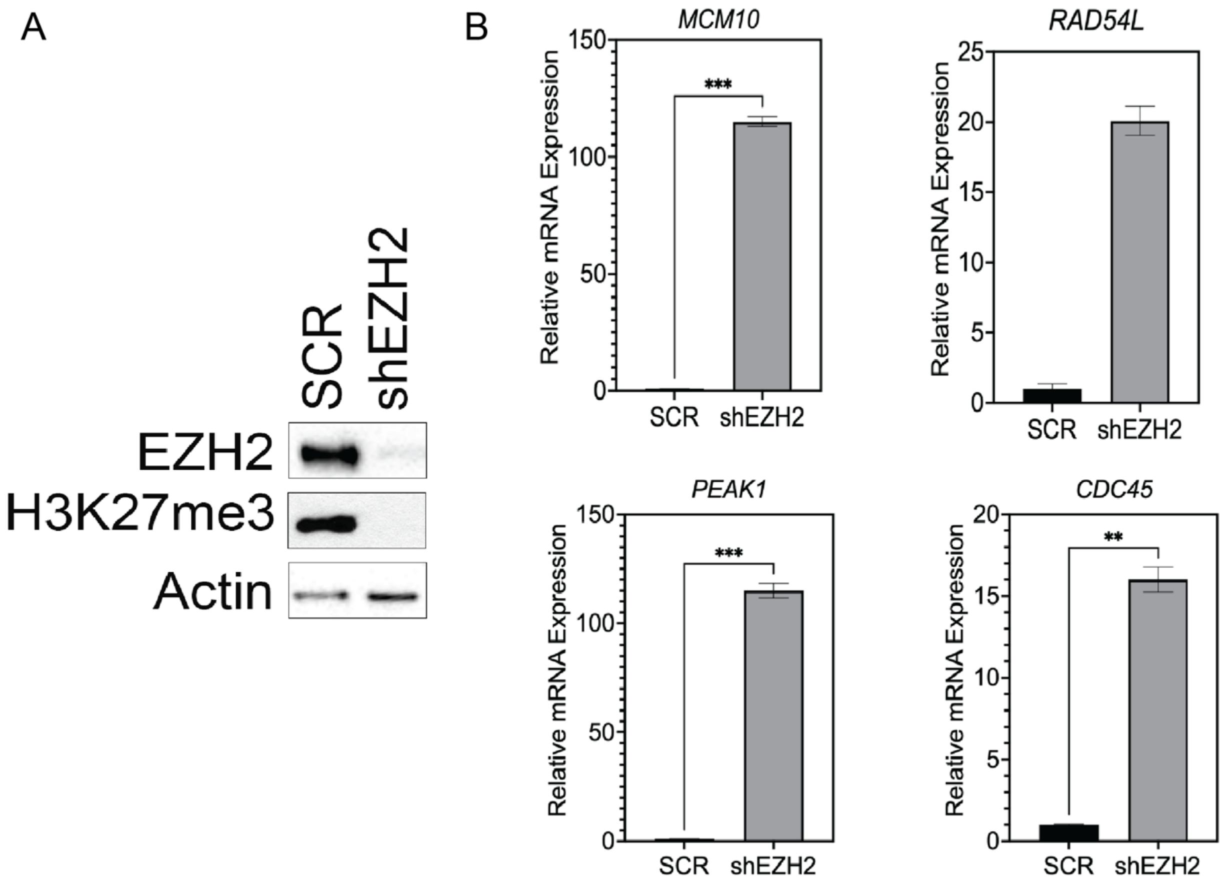


**Fig. S1. Validation of EZH2 Target genes.**

Genes significantly enriched in the retrospective RNA-seq analysis were confirmed as "true" PRC2 targets via RT-qPCR in 1833 cells after knockdown of primary catalytic component of the complex, EZH2. Gene expression was normalized to a housekeeping gene (18s) before being calculated relative to SCR transduced cells. Significantly enriched genes are marked as indicated. Data are representative of 3 independent experiments.

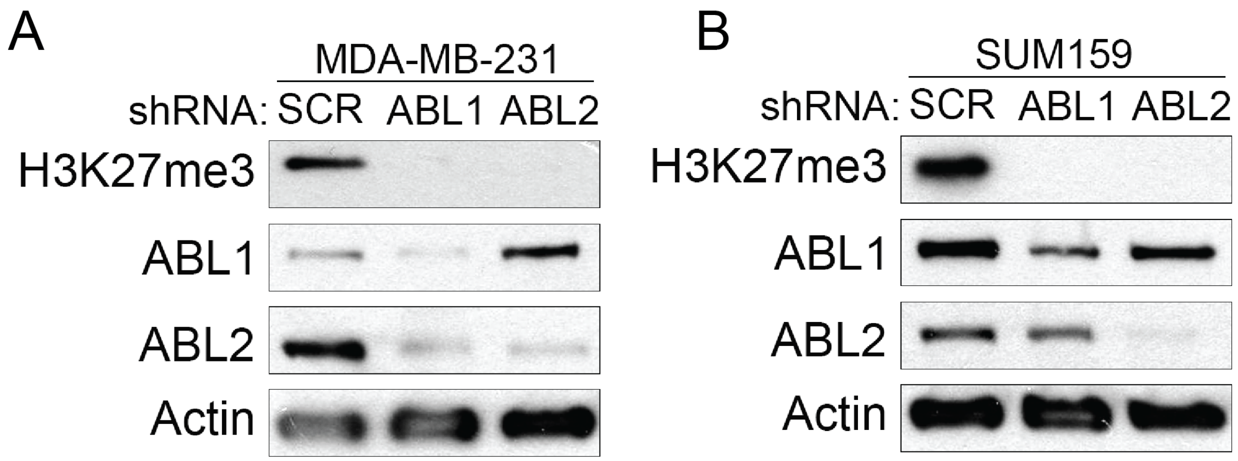


**Fig. S2. Single ABL Knockdown ablates H3K27me3.** MDA-MB-231 cells were transfected with either SCR nontargeting shRNA or shRNAs against ABL1 or ABL2 respectively. Cell lysates were prepared and analyzed by immunoblotting for H3K27me3 levels. Actin was used as a loading control. Data are representative of 3 independent experiments.

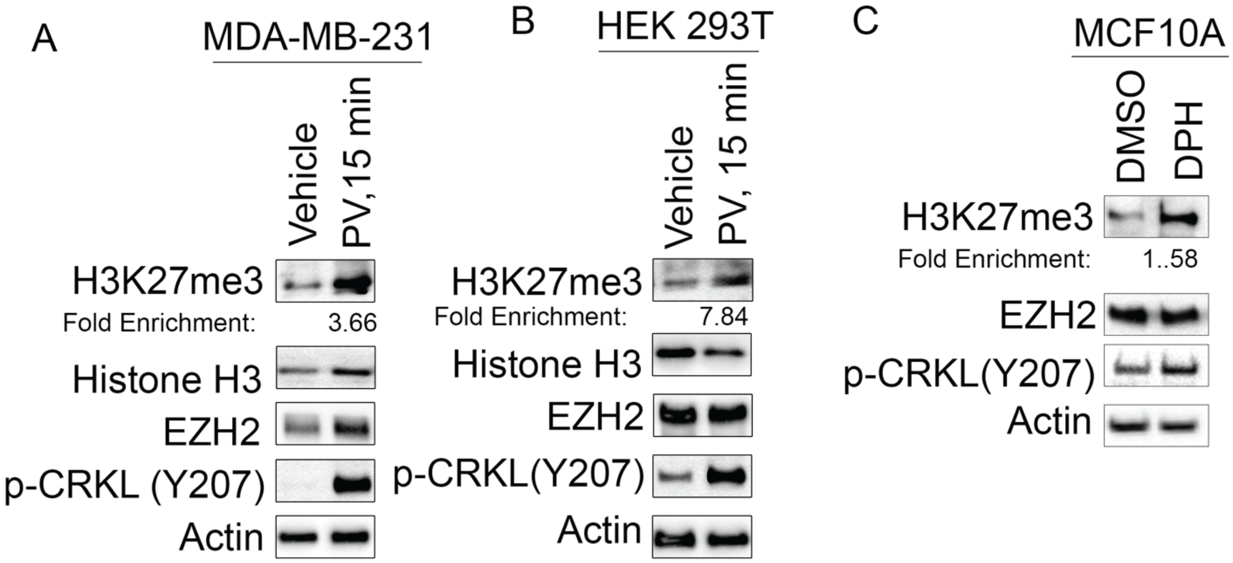


**Fig. S3. Tyrosine Phosphorylation Regulates EZH2 catalytic activity.** Cells were treated with 50uM Pervanadate (PV) for 15 minutes, after which cells were assessed via immunoblot. Actin was used as a loading control. Inhibition of tyrosine phosphatases is sufficient to induce H3K27me3 levels in (A) MDA-MB-231 and (B.) HEK 293T Cells. (C.) Untransformed mammary epithelial cells (MCF10A) were treated 10uM DPH for 24 hours cells were then assessed via immunoblot for changes in H3K27me3 levels. Actin was used as a loading control. Data are representative of 3 independent experiments.

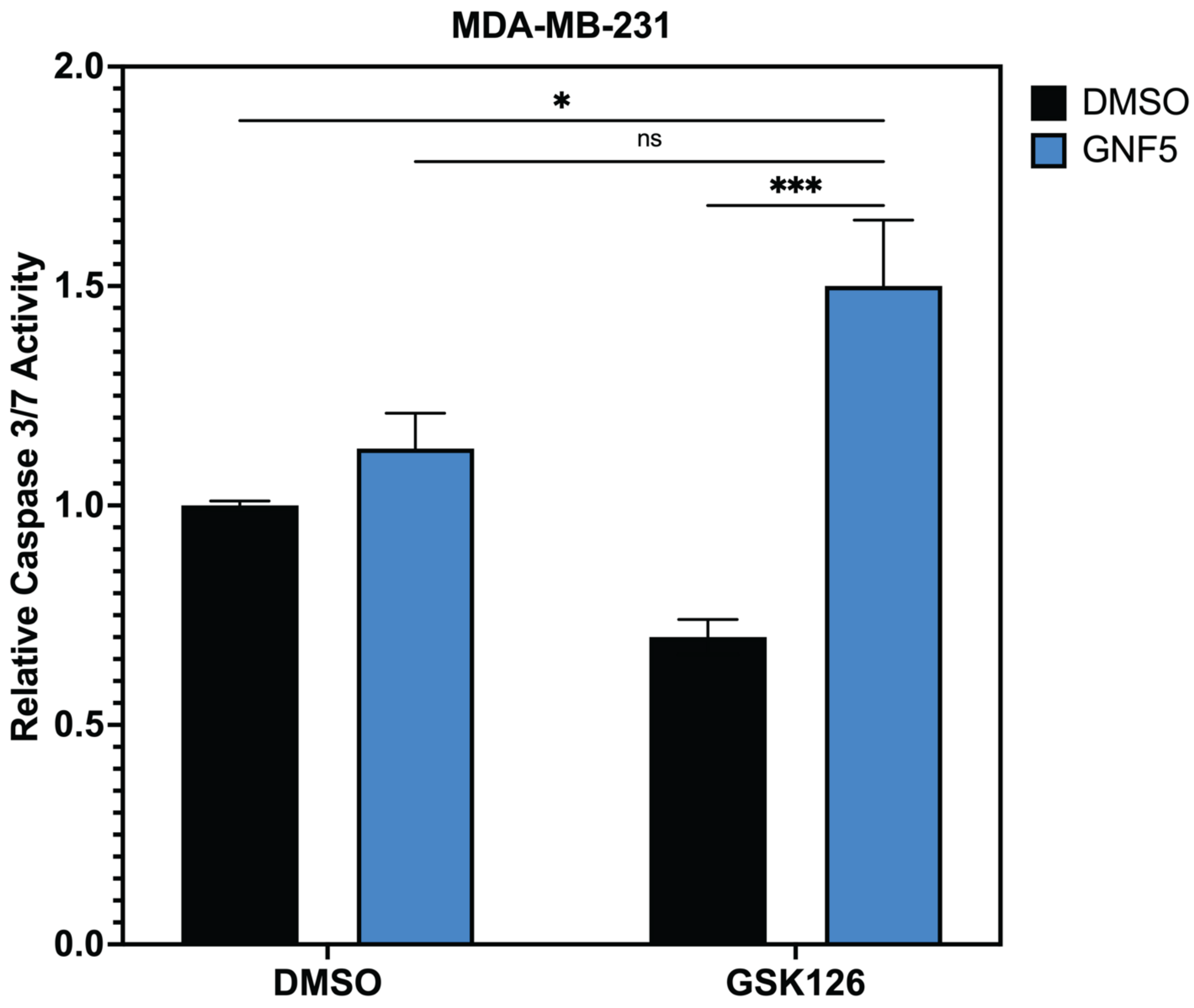


**Fig. S4. Combination treatment with ABL inhibitors and EZH2 inhibitors induces caspase activity.**

MDA-MB-231 cells were treated with either DMSO, 10uM GNF5, 10uM GSK126, or both GNF5 and GSK126 for 24 hours after which Caspase 3/7 activity was assessed using Caspase Glo. Caspase activity was normalized to DMSO treated cells. Data represent results of 3 independent experiments. Significance between treatment groups was determined using a two-way analysis of variance followed by Tukey's post hoc testing, asterisks indicate significance, *,$p<0.05$, ns=not significant. Data are representative of 3 independent experiments.

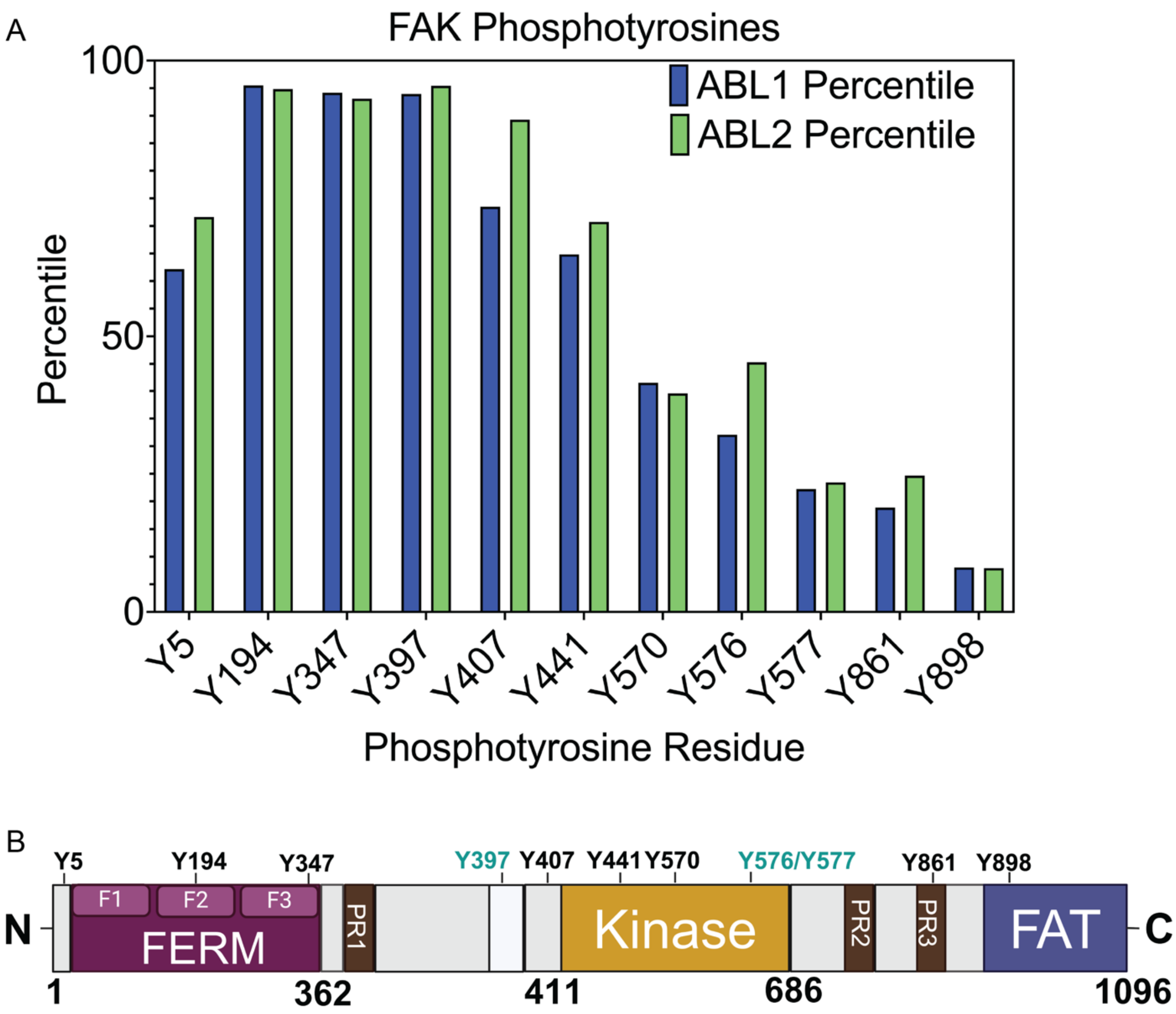


**Fig. S5. FAK is a putative ABL Kinase Target.**
(A.) Predicted phosphorylation sites were extrapolated from data on phosphosite. (B.) Linear protein structure of FAK with predicted phosphosites marked as indicated. Key functional residues are highlighted in green.

**Table S1.** CDK1 Phosphotyrosine Sites in Invasive Breast Carcinoma

| Residue | Post Translational Modification | Site | Predicted Kinase |
|---|---|---|---|
| **Tyrosine** | Phosphorylation | y15 | FYN, SRC,HCK,LYN, BLK, **FAK** |
| **Tyrosine** | Phosphorylation | y15y19 | WEE1 |
| **Tyrosine** | Phosphorylation | y160 | FES, **FAK** |
| **Tyrosine** | Phosphorylation | y19 | WEE1 |
| **Tyrosine** | Phosphorylation | y286 | FGFR1, FGFR3,TYK2 |

**Table S2**. RT-qPCR Primers

| Gene | Forward Primer | Reverse Primer |
|---|---|---|
| 18s | 5’- GAGGATGAGGTGGAACGTGT-3’ | 5'- AGAAGTGACGCAGCCCTCTA-3’ |
| CDC45 | 5’-TTCGTGTCCGATTTCCGCAAA-3’ | 5’- TGGAACCAGCGTATATTGCAC-3’ |
| MCM10 | 5’- TGTCCCTGCGCTACCAAGA- 3’ | 5’- GATGAGCTTTTGGGATCTGGAG-3’ |
| RAD54L | 5’- AGGCAGGTCCTGTGATGATGA-3’ | 5’- TCAAAGGTTTCCGAAAAGGAGAC-3’ |
| PEAK1 | 5’- TAGTGGCAGATGGGCAAAGTA-3’ | 5’- TGTTTCGGTTCCACCCTATGA-3’ |

**Table S3.** shRNAs

| Construct | Sequence | Backbone | Source |
|---|---|---|---|
| SCR | GGTGTATGGGCTACTATAGAA | pLKO.1 | Previously described in (36) |
| shABL1 | GGTGTATGAGCTGCTAGAGAA | pLKO.1 | |
| shABL2 | CCTTATCTCACCCACTCTGAA | pLKO.1 | |
| shAA | AGGTACTAAAGTGGCTCTGAG | pLKO.1 | |
| shEED #1 | CCGGGCAAACTTTATGTTTGGGATTCTCGAG AATCCCAAACATAAAGTTTGCTTTTT | pLKO.1 | Duke Functional Genomics Shared Resource Facility |
| shEED #2 | CCGGCCAGAGACATACATAGGAATTCTCGAG AATTCCTATGTATGTCTCTGGTTTTT | pLKO.1 | Duke Functional Genomics Shared Resource Facility |
| shEZH2 | CCGGCAACACAAGTCATCCCATTAACTCGAG TTAATGGGATGACTTGTGTTGTTTTTG | pLKO.1 | Duke Functional Genomics Shared Resource Facility |
| shFAK #1 | CCGGGAGAGCATGAAGCAAAGAATTCTCGA GAATTCTTTGCTTCATGCTCTCTTTTT | pLKO.1 | Duke Functional Genomics Shared Resource Facility |
| shFAK #2 | CCGGCGGAGAATATGGCTGACCTAACTCGAG TTAGGTCAGCCATATTCTCCGTTTTT | pLKO.1 | Duke Functional Genomics Shared Resource Facility |

**Table S4.** Antibodies

| Antibody | Manufacturer | Catalogue Number | RRID |
|---|---|---|---|
| ABL1 | BD Biosciences | 554148 | AB_2220994 |
| ABL2 | Abnova | H00000027-M03 | AB_565433 |
| β-Actin | Cell Signaling Technology | 3700 | AB_2242334 |
| CDK1 | Cell Signaling Technology | 9116 | AB_2074795 |
| Phospho-CDK1 Y15 | Cell Signaling Technology | 4539 | AB_560953 |
| Phospho-CDK1 T161 | Cell Signaling Technology | 9114 | AB_2074652 |
| CRKL | Santa Cruz Biotechnology | sc-365092 | AB_10847682 |
| Phospho-CRKL | Cell Signaling Technology | 3181 | AB_331068 |
| EED | Cell Signaling Technology | 85322 | AB_2923355 |
| EZH2 | Cell Signaling Technology | 5246 | AB_10694683 |
| Phospho-EZH2 T487 | Thermo Fisher Scientific | PA5105660 | AB_2817088 |
| FAK | Cell Signaling Technology | 71433 | AB_2799801 |
| Phospho-FAK Y397 | Cell Signaling Technology | 3283 | AB_2173659 |
| Monoclonal Anti-Flag® M2 | Millipore Sigma | F1804 | AB_262044 |
| GAPDH | Santa Cruz Biotechnology | sc-32233 | AB_627679 |
| GFP | Cell Signaling Technology | 2956 | AB_1196615 |
| HA | Cell Signaling Technology | 3724 | AB_1549585 |
| Histone H3 | Cell Signaling Technology | 4499 | AB_10544537 |
| H3K27me3 | Cell Signaling Technology | 9733 | AB_2616029 |
| c-MYC | Cell Signaling Technology | 5605 | AB_1903938 |
| p-Y 4G10 | Millipore Sigma | 05-321 | AB_2891016 |
| SUZ12 | Cell Signaling Technology | 3737 | AB_2196850 |
| β-Tubulin | Cell Signaling Technology | 2146 | AB_2210545 |
| Vinculin | Cell Signaling Technology | 4650 | AB_10559207 |
| ZMYND8 | Cell Signaling Technology | 97845 | AB_2800291 |
